# Mechanism of GPR84 allosteric modulation at a helix 8-proximate site

**DOI:** 10.64898/2026.04.10.715585

**Authors:** Xuan Zhang, Abdul-Akim Guseinov, Laura Jenkins, Jingkai Zhou, Fabian Gossen, Pinqi Wang, Zobaer Al Mahmud, Yueming Li, Andhika B. Mahardhika, Christa E. Müller, Mingye Feng, Angela J. Russell, Irina G. Tikhonova, Graeme Milligan, Cheng Zhang

## Abstract

Allosteric modulators offer opportunities for pathway-selective GPCR signalling, but the structural mechanisms enabling biased allosteric modulation remain unclear. Here we identify a helix 8-proximate allosteric site in the immune-metabolic receptor GPR84 and define how it achieves G_i_ -biased signalling. Cryo-EM structures of the GPR84-G_i_ complexes bound to the orthosteric agonist OX04539 alone or in combination with the positive allosteric modulator (PAM) PSB-16671 reveal that PSB-16671 binds at the interface of TM1, TM7, and helix 8, a location distinct from previously characterized GPCR allosteric pockets. Molecular dynamics simulations and mutagenesis uncover a polar interaction network linking orthosteric and allosteric sites through conserved residues including Asp66^2.50^, Asn104^3.36^, and Asn362^7.45^. Unexpectedly, disrupting this network enhances allosteric cooperativity, indicating that conformational flexibility within the network is essential for allosteric communication. PSB-16671 stabilizes a receptor conformation with pronounced TM6 displacement that favours Gi coupling while disfavouring β-arrestin recruitment. This G_i_-biased profile sustains macrophage phagocytosis of cancer cells without the desensitization induced by balanced agonists. Sequence analysis suggests that helix 8-proximate allosteric sites may be broadly targetable across class A GPCRs, while receptor-specific contacts enable selective modulation. These findings establish structural and mechanistic principles for biased allosteric modulation applicable beyond GPR84.

## Introduction

GPR84 is a family A G protein-coupled receptor (GPCR) predominantly expressed in innate immune cells such as neutrophils, monocytes, and macrophages^1–3^. Initially identified as a receptor for medium-chain fatty acids (MCFAs), GPR84 is now recognized as an immune-metabolic receptor that regulates innate immune responses^1,3–6^. Its expression is rapidly upregulated by pro-inflammatory stimuli, including lipopolysaccharide (LPS) and interferon-γ (IFN-γ), particularly in activated microglia and macrophages^2,3,7^. Functionally, GPR84 signaling amplifies inflammatory responses, positioning it as a potential driver of various immune-mediated pathologies such as metabolic syndrome, fibrosis, inflammatory bowel disease, and neuroinflammation^3,4^. As a result, blocking GPR84 signaling represents a therapeutic strategy for these disorders. The small-molecule antagonist, GLPG1205 (9-(2-cyclopropylethynyl)-2-[[(2*S*)-1,4-dioxan-2-yl]methoxy]-6,7-dihydropyrimido[6,1-*a*]isoquinolin-4-one), showed promise in preclinical models of idiopathic pulmonary fibrosis (IPF) and ulcerative colitis (UC)^3,8,9^, although it failed to meet efficacy endpoints in later-stage clinical trials^10^. Beyond inflammation, GPR84 regulates macrophage phagocytosis^2,11^ with activation by the orthosteric agonist 6-OAU (6-(octylamino)pyrimidine-2,4(1*H*,3*H*)-dione) promoting macrophage-mediated phagocytosis of cancer cells^1,5,12^, highlighting potential applications in cancer immunotherapy.

Despite early identification of endogenous and surrogate agonists, including decanoic acid (capric acid, C10) and 2-(hexylthio)pyrimidine-4,6-diol (**Fig. 1a**)^4,6^, these ligands exhibit limited potency, metabolic stability, or receptor selectivity, restricting their utility as pharmacological tools or therapeutic leads. Recent synthetic efforts have yielded orthosteric agonists with improved potency and drug-like properties^13,14^. In addition, identification of 3,3′-diindolylmethane (DIM) (**Fig. 1a**), a compound derived from cruciferous vegetables, as both a GPR84 activator^15^ and positive allosteric modulator (PAM)^16^ showed that allosteric pockets exist on GPR84 and provide alternative regulatory mechanisms. Structure-activity relationship (SAR) studies^16^ led to the development of more potent DIM analogs, notably PSB-15160 (5-fluoro-3-[(5-fluoro-1*H*-indol-3-yl)methyl]-1*H*-indole) and PSB-16671 (3-[(5,7-difluoro-1*H*-indol-3-yl)methyl]-5,7-difluoro-1*H*-indole)^16^ (**Fig. 1a**).

**Fig. 1.**
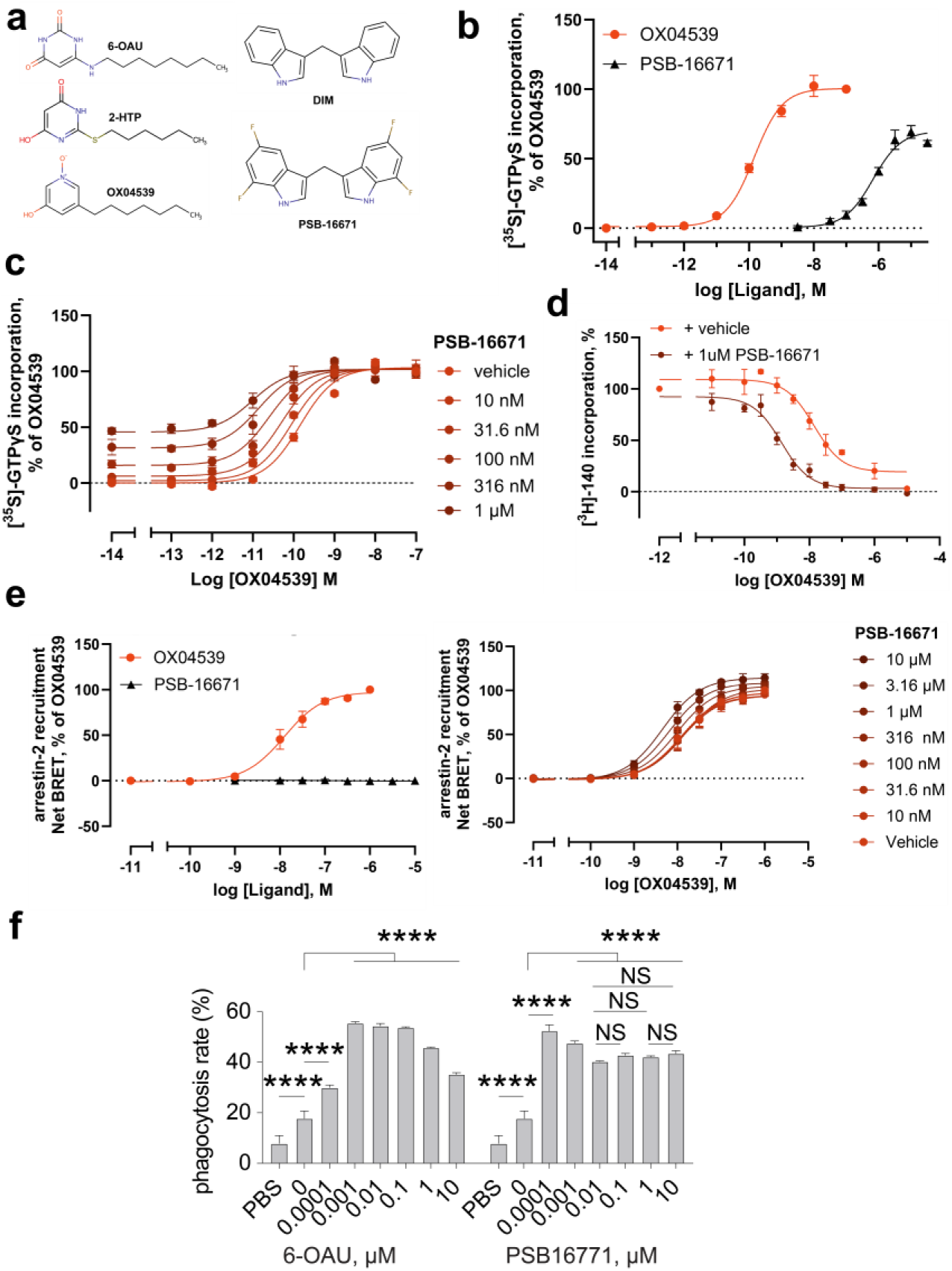
Pharmacological characterization of PSB-16671 as a G_i_-biased ago-PAM of GPR84. **a.** Chemical structures of the three orthosteric ligands 6-OAU, 2-HTP, and OX04539, and the two allosteric modulators DIM and PSB-16671**. b**. Both PSB-16671 and OX04539 directly promote binding of [^35^S]GTPγS to a human GPR84-G_i_ fusion protein in a concentration-dependent manner. **c.** Increasing concentrations of PSB-16671 increase the potency of OX04539 to enhance binding of [^35^S]GTPγS to the GPR84-G_i_ fusion protein. **d.** The potency of OX04539 to compete with [^3^H]140 to bind GPR84 is increased in the presence of 1µM PSB-16671. **e.** PSB-16671 increases the potency of OX04539 to promote interactions between GPR84 and arrestin-2 without PSB-16671 directly promoting such an interaction. All data represent means ± SEM, n = 3 independent experiments for **b-e**. **f.** Dose-dependent pro-phagocytic effect of 6-OAU and PSB-16671. All data points except for the PBS data were obtained in the presence of anti-CD47 antibodies. PSB-16671 promotes sustained cancer cell phagocytosis, while high concentrations of 6-OAU cause desensitization. Each data point represents SD of data from 3 independent experiments (n=3). NS, not significant. *\*\*\*\*p<0.001*.

We previously determined a high-resolution cryo-EM structure of a GPR84–G_i_ complex bound to 6-OAU (PDB ID 8G05)^5^ and additional structures with different orthosteric agonists have been reported^17,18^. However, the binding site for allosteric modulators including DIM and PSB-16671 remained unknown. Here we define this site using a cryo-EM structure of the GPR84-G_i_ complex bound to both PSB-16671 and the potent orthosteric agonist OX04539 (**Fig. 1a**)^14^, revealing an allosteric site not previously reported in family A GPCRs. We show that PSB-16671 produces sustained enhancement of macrophage phagocytosis and, through integrated approaches combining structural biology, molecular dynamics (MD) simulations, mutagenesis and medicinal chemistry, elucidate the molecular mechanisms underlying the allosteric modulation and biased signaling of PSB-16671.

## Results

### PSB-16671 is a G_i_-biased ago-PAM of GPR84

To characterize the function of PSB-16671 as a direct activator of GPR84-G_i_-signaling we conducted [^35^S]GTPγS binding studies using membranes of Flp-In T-REx 293 cells induced to express a human GPR84-G_i2_α fusion protein^19^. PSB-16671 produced a concentration-dependent increase in binding of [^35^S]GTPγS with moderate potency (**Fig. 1b**). In comparison, the recently described GPR84 orthosteric agonist OX04539^14^ was both more potent and displayed greater efficacy than PSB-16671 (**Fig. 1b**, **Table 1**). Co-addition studies showed co-operativity between the two ligands, with PSB-16671 acting as a positive allosteric modulator (PAM) for G_i_-activation by OX04539 (**Fig. 1c**), primarily by increasing the potency of OX04539 (**Fig. 1c)**. This implies that PSB-16671 and OX04539 must bind to different sites on GPR84. Analysis of such data using the operational model of allosterism predicted the affinity (logK_d_) of PSB-16671 to be -6.20 ± 0.23 and that of OX04539 -8.62 ± 0.20 (means ± SD, n = 3). As anticipated from the functional co-operativity, co-addition of PSB-16671 (1µM) to experiments that measured competition between OX04539 and the GPR84 orthosteric antagonist [^3^H]140^20,21^ to bind to GPR84 demonstrated increased affinity of OX04539 in the presence of PSB-16671 (calculated logK_d_ = -9.21 ± 0.14 compared to -8.20 ± 0.32 in the absence of PSB-16671, means ± SD, n = 3) (**Fig. 1d**), and that PSB-16671 had very limited direct effect on the binding of [^3^H]140.

**Table 1.** Potency and efficacy of orthosteric and allosteric agonists at wild type and GPR84 Thr^41^Val.

|  | **logEC<sub>50</sub>, M** | **Top, % of OX04539** |
| --- | --- | --- |
| **OX04539** | -9.83 ± 0.07 | 102 ± 8 |
| **2-HTP** | -7.67 ± 0.06 | 101 ± 8 |
| **PSB-16671** | -6.17 ± 0.13 | 70 ± 6 |
| **PSB-16242** | N.D. | N.D. |
| **PSB-16244** | -5.18 ± 0.10*** | 59 ± 16 |
| **PSB-2502** | -5.59 ± 0.24** | 84 ± 18 |
| **PSB-2504** | -5.81 ± 0.22 | 74 ± 22 |
| **PSB-2505** | -6.00 ± 0.15 | 81 ± 22 |
| **DIM** | N.D. | N.D. |
| **Thr<sup>41</sup>Val, OX04539** | -10.14 ± 0.09 | 97 ± 2 |
| **Thr<sup>41</sup>Val, 2-HTP** | -8.09 ± 0.09 | 98 ± 3 |

Previous studies have characterized PSB-16671 as a G protein-biased ligand, showing, unlike OX04539^18^ and 6-OAU, an inability to directly promote arrestin interactions with GPR84^4,6,22^. We confirmed this (**Fig. 1e**). However, in co-addition studies, PSB-16671 was able, in a concentration-dependent manner, to enhance the potency of OX04539 to recruit arrestin-2 to GPR84 (**Fig. 1f)**, consistent with the increased affinity of OX04539 in the presence of PSB-16671 as detected in [^3^H]140/OX04539 competition binding assays. This indicates that binding of PSB-16671 does not preclude arrestin interaction with GPR84. Instead, it likely induces specific receptor conformations sufficient for the coupling of G_i_ but not direct arrestin recruitment.

Nevertheless, we reasoned that the G_i_-biased property of PSB-16671 may lead to less receptor desensitization compared to balanced agonists such as 6-OAU. In a previous study we demonstrated that 6-OAU stimulates GPR84-G_i_ signaling in macrophages to promote cancer cell phagocytosis^5^. However, high 6-OAU concentrations led to reduced phagocytosis (**Fig. 1f**), likely due to GPR84 desensitization^14^. In contrast, PSB-16671 maintains phagocytic activity across a large concentration range without apparent desensitization (**Fig. 1f**), consistent with the G_i_-biased property of this ago-PAM.

### Overall structure of the GPR84-G_i_ signaling complex with OX04539 and PSB-16671

To elucidate the molecular basis for the PAM effect of PSB-16671 on function of OX04539, we obtained cryo-EM structures of human GPR84 in complex with a heterotrimeric G_i_ protein bound to OX04539 alone or in combination with PSB-16671 at overall resolutions of 3.43 Å and 3.22 Å, respectively (**Fig. 2a, Fig. S1-2, Table. S2**). The structure determination largely followed our previous methods for obtaining the structure of the GPR84-G_i_ complex with 6-OAU^5^. For the GPR84-G_i_ complex with both ligands, we added PSB-16671 to the complex purified with OX04539 before making the cryo-EM grids.

**Fig. 2.**
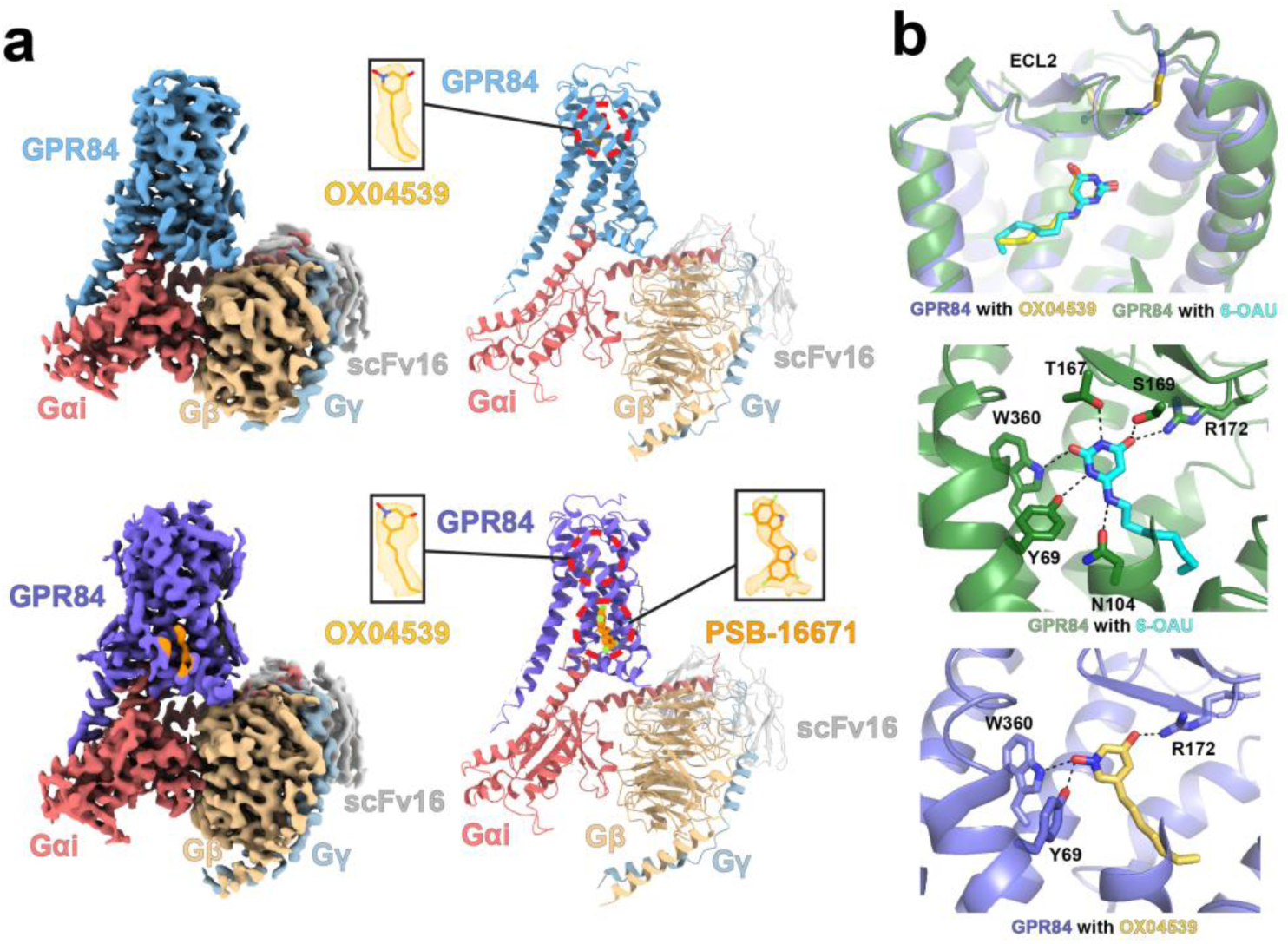
Structures of the GPR84-G_i_ signaling complex with OX04539 and PSB-16671. **a.** Cryo-EM density maps and overall structures of the GPR84-G_i_ complex with OX04539 alone (upper) and both OX04539 and PSB-16671 (lower). GPR84 is colored in deep teal and slate, respectively, in the complexes with OX04539 alone and with both ligands. G_αi_, Gβ and Gγ subunits are colored in red, sand and light blue, respectively. ScFv16 is colored in grey. The cryo-EM density maps of ligands are shown as meshes. **b.** Binding poses of OX04539 and 6-OAU in the orthosteric binding pocket. Polar interactions are shown as black dashes lines.

The cryo-EM maps revealed clear density for OX04539, which closely resembles the binding pose of 6-OAU^5^ in such a complex (**Fig. 2b**). Nearly all residues within the orthosteric binding pocket, including those from extracellular loops (ECLs), adopt similar conformations in each of these structures, indicating a conserved binding mechanism for each orthosteric ligand. However, since the head group of OX04539 contains fewer amine groups (**Fig. 1a**), it forms polar interactions only with Tyr69^2.53^, Arg172^ECL2^, and Trp360^7.43^ (superscripts represent Ballesteros-Weinstein numbering ^23,24^), whereas 6-OAU engages in more extensive polar interactions with the receptor (**Fig. 2b**). On the other hand, the tail group of OX04539 adopts a more extended conformation, while the longer tail group of 6-OAU is bent at its terminus due to steric constraints (**Fig. 2b**), which may lead to lower potency of 6-OAU compared to OX04539.

In the case of PSB-16671 we observed additional cryo-EM density located at the interface between transmembrane helices 1 and 7 (TM1 and TM7) and helix 8, which well accommodates the ligand (**Fig. 2a**). Such density was absent in the cryo-EM map of the GPR84-G_i_ complex with OX04539 alone (**Fig. 3a**), confirming the distinct allosteric binding site for PSB-16671. Further validation will be discussed in later sections.

**Fig. 3.**
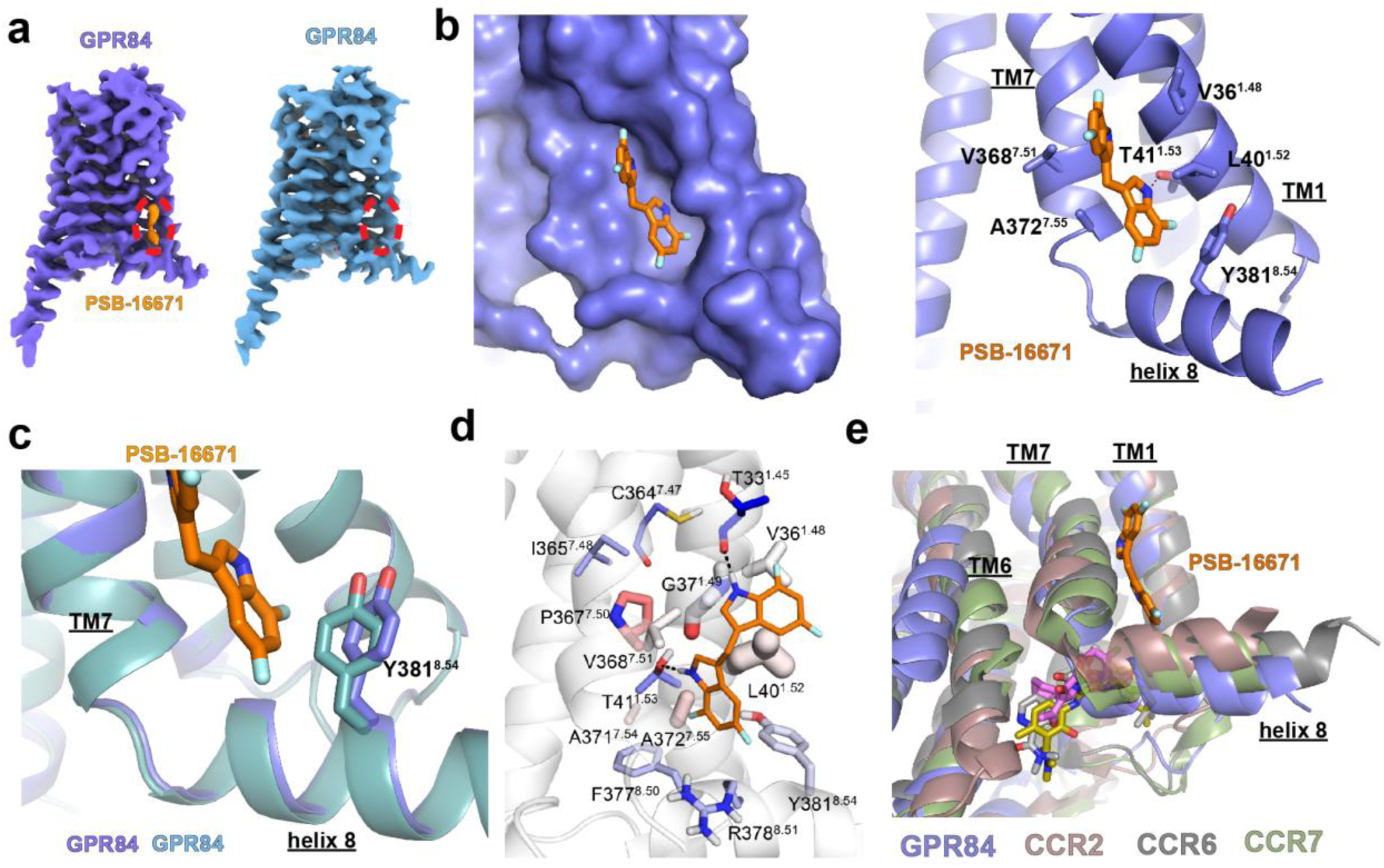
Binding of PSB-16671 at a helix 8-proximate allosteric site. **a.** Density of PSB-16671 (orange) present in the cryo-EM map of the GPR84-G_i_ complex with both OX04539 and PSB-16671 (slate) but not in the map of the complex with OX04539 alone (deep teal) **b.** Binding of PSB-16671 at the allosteric site. **c.** Different conformation of the Tyr381^8.54^ side chain in GPR84 with (slate) and without (deep teal) PSB-16671. **d.** Simulation snapshot of the allosteric site. Residue colour and thickness represent the calculated energies of electrostatic and van der Waals interactions, respectively. Polar interactions are shown as black dashes lines in **b** and **d**. **e.** Structural alignment of GPR84 (slate), CCR2 (brown), CCR6 (dark gray), and CCR7 (green) with intracellular allosteric modulators.

### Helix 8-proximate allosteric site of PSB-16671

PSB-16671 exhibits a symmetric structure with two identical difluoro-indole rings connected by a methylene bridge (**Fig. 1a**). It sits in a pocket formed among TM1, TM7, and helix 8, adopting an extended conformation along these transmembrane helices (**Fig. 3b**). One indole ring engages in a hydrogen bond with Thr41^1.53^ of GPR84 through its NH group, while forming hydrophobic and π-π interactions with GPR84 residues Leu40^1.52^, Ala372^7.55^, and Tyr381^8.54^ (**Fig. 3b**). The other indole ring makes hydrophobic interactions with Val36^1.48^ and Val368^7.51^ from TM1 and TM7 of GPR84, respectively (**Fig. 3b**). Notably, PSB-16671 binding induces a conformational shift of the Tyr381^8.54^ side chain away from TM7 compared to its position in the GPR84-G_i_ complex with OX04539 alone due to steric effects (**Fig. 3c**).

Molecular dynamics (MD) simulations validated the structural stability of this complex, confirming the cryo-EM binding mode of PSB-16671 (**Table S3**). As expected from its lower estimated binding affinity, PSB-16671 exhibited reduced stability compared to the orthosteric ligand OX04539. The simulations confirmed a persistent hydrogen bond between one indole ring of PSB-16671 and Thr41^1.53^, while revealing that the second indole ring forms a hydrogen bond predominantly with the backbone oxygen of Thr33^1.45^ and, to a lesser extent, with the backbone of Cys364^7.47^ (**Fig. 3d, Fig. S3a**). Residue-ligand interaction energy calculations identified strong van der Waals interactions with Leu40^1.52^, Gly37^1.49^, Ala372^7.55^, Val36^1.48^, and Val368^7.51^, along with notable electrostatic and van der Waals contributions from Arg378^8.51^, Tyr381^8.54^, and Phe377^8.50^ (**Fig. 3d, Table S4**).

The location of the binding site of PSB-16671, near helix 8 of the receptor and lipid interface, was unanticipated. Intracellular allosteric sites involving helix 8 have been reported for various chemokine receptors including CCR2, CCR6, and CCR7^25–27^, where negative allosteric modulators (NAMs) bind and interact with TM1, TM6, and helix 8 (**Fig. 3e**). Due to a steric clash, those NAMs block G protein coupling directly. These sites differ from the PSB-16671 site in both location and mechanism of allosteric modulation (**Fig. 3e**). In contrast to NAMs that sterically block G-protein engagement, PSB-16671 appears to stabilize TM7 and helix 8 in an active conformation that enhances G-protein coupling, which is elaborated in a later section.

### Mutagenesis and structure-activity relationship validation of PSB-16671 biding mode

The identified allosteric binding site of PSB-16671 and the molecular interactions with the receptor, although unanticipated, are consistent with previous structure-activity relationship (SAR) data, which demonstrates the critical importance of both indole NH groups as hydrogen bond donors. Methylation of one or both indole nitrogens completely eliminates GPR84 activity^14,16^, confirming the essential role of these polar interactions observed in the structure and MD simulations. In addition, introduction of any alkyl or aryl substituents at the methylene bridge completely abolishes activity, with oxidation to a carbonyl being the only modification that preserves partial activity (approximately 2-fold reduced potency)^14,16,28^. This strict structural requirement aligns with the location of the bridge at the confined interface between TM1, TM7, and helix 8, where steric constraints are severe and the dynamic flexibility of the CH2 group appears crucial for optimal positioning of both indole rings within the binding pocket. The steric sensitivity of this binding site is also demonstrated by the SAR of symmetrically substituted PSB-16671 analogs^14,16,28^. Only small fluorine substituents are well-tolerated at the 5,7-positions, while bulky groups or strong electron-withdrawing substituents dramatically reduce activity^14,16,28^.

To further validate the PSB-16671 binding site revealed by our cryo-EM structure, which showed a critical hydrogen bond between PSB-16671 and Thr41^1.53^ of GPR84 as representing the primary polar interaction within the binding pocket (**Fig. 3b**), we performed a series of mutagenesis and signaling experiments. In addition, to assess potential differential steric requirements of each indole ring, we explored unsymmetrical analogs, including PSB-16242 with N-methylation predicted to prevent hydrogen bonding, and PSB-16244 with a single 5-fluorine substitution ^16,28^ (**Fig. 4a**). We additionally synthesized (**Scheme 1**) three further compounds PSB-2502, PSB-2504, and PSB-2505 (**Fig. 4a**) to test whether fluorination of one of the indole rings is more important than the other. Functional evaluation using [^35^S]GTPγS binding studies at the GPR84-G_i2_α fusion protein revealed that, as anticipated, PSB-16242 was almost inactive (**Fig. 4b**), confirming that N-methylation disrupts essential hydrogen bonding. By contrast, unsymmetrical modifications were well-tolerated, with PSB-2502, PSB-2504, and PSB-2505 showing similar potencies (pEC_50_ = -5.59-6.00) regardless of fluorine positioning (**Fig. 4b**, **Table 1**). Alteration of Thr41^1.53^ to Val was predicted to prevent the hydrogen bond to the indole NH and indeed PSB-16671 was unable to stimulate [^35^S]GTPγS binding to a GPR84 Thr41^1.53^Val-G_i2_α fusion protein (**Fig. 4c**). Moreover, the GPR84 Thr41^1.53^Val mutation eliminated response to DIM and all the PSB-16671 analogs tested (**Fig. 4c)**. Importantly, however the Thr41^1.53^Val mutation preserved response to OX04539, as well as to a second orthosteric agonist 2-HTP (also designated as ZQ-16^15,22^) (**Fig. 4b and c**). Additionally, this mutation did not alter the binding affinity of the GPR84 orthosteric antagonist [^3^H]140 (K_d_ of wt GPR84 is 4.74 ± 1.84 nM and of Thr41^1.53^Val is 3.97 ± 1.07 nM, means ± SD, n = 3) (**Fig. 4d**). This confirms that the Thr41^1.53^Val mutation neither prevented expression of a functional GPR84 receptor nor abolished the intrinsic signaling capability of GPR84. Furthermore, there was a lack of co-operative effect of PSB-16671 on the function of OX04539 at GPR84 Thr41^1.53^Val in the [^35^S]GTPγS binding assay (**Fig. 4e**) and PSB-16671 was unable to increase the observed affinity of OX04539 at GPR84 Thr41^1.53^Val (logK_d_ = - 8.20 ± 0.05, mean ± SD. n = 3) in [^3^H]140 competition binding studies (**Fig. 4f**).

**Fig. 4.**
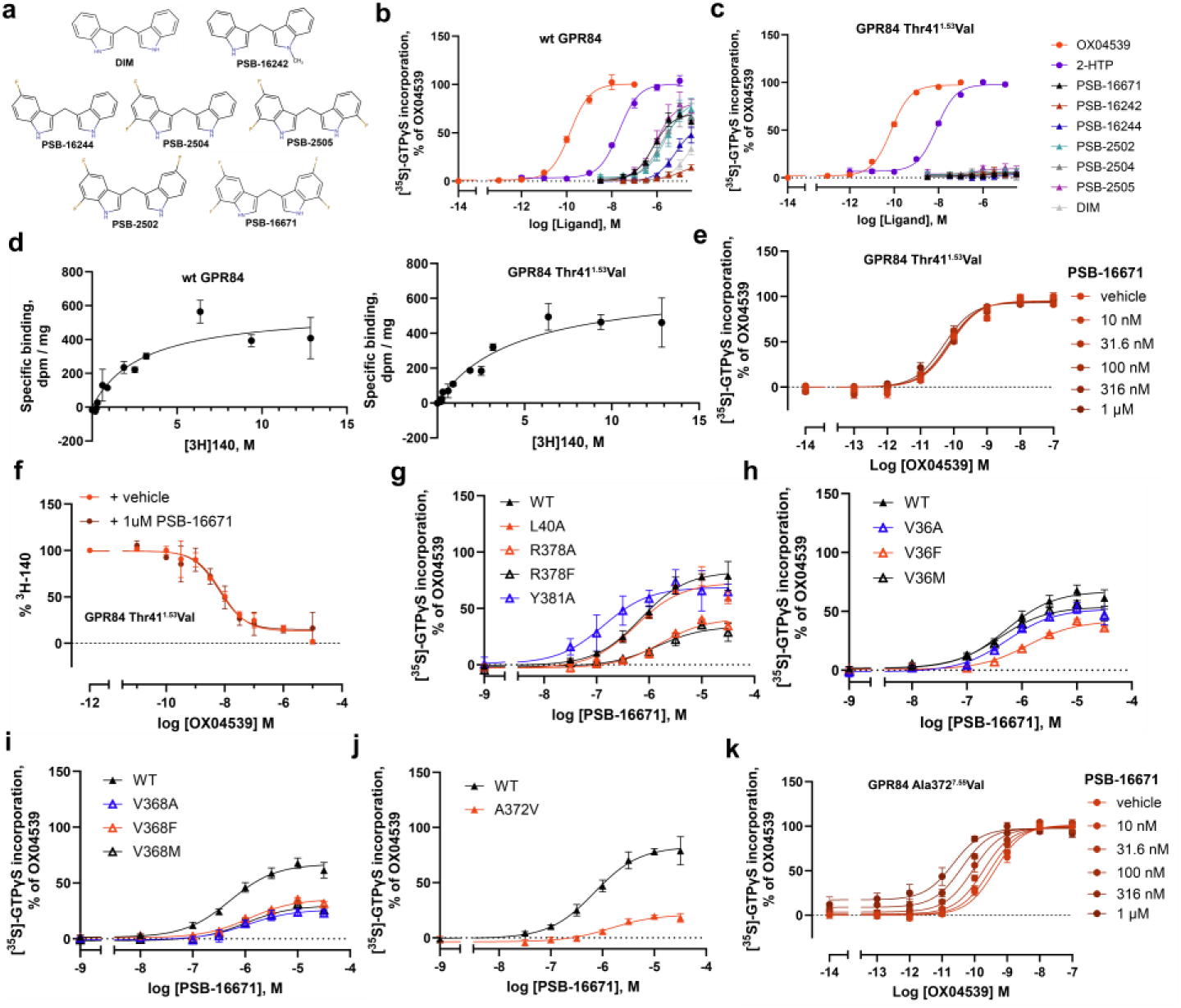
Functional validation of PSB-16671 binding site. **a**. Chemical structures of analogs of PSB-16671. **b**. PSB-16242 lacks significant potency at wild type GPR84 but fluorinated analogs, PSB-16244, PSB-2502, PSB-2504 and PSB-2505, display similar potency in GTPγS-binding assays. **c**. All analogs of PSB-16671 fail to directly activate G_i_ via GPR84 Thr41^1.53^Val whilst the orthosteric agonists OX04539 and 2-HTP do. **d**. The orthosteric GPR84 antagonist [^3^H]140 binds wild type and GPR84 Thr41^1.53^Val with similar affinity. **e.** PSB-16671 lacks co-operative activity with OX04539 at GPR84 Thr41^1.53^Val at concentrations up to 1μM and, **f**. does not alter the affinity of OX04539 to compete with [^3^H]140 to bind GPR84. **g-j.** Effects of mutations at the allosteric binding site on the agonist activity of PSB-16671. **k.** The Ala372^7.55^Val mutation does not reduce the extent of allosteric co-operativity between OX04539 and PSB-16671. Quantitative analysis is shown in **Table 2**. Each data point in **b-k** represents Means +/- S.E.M. from 3 independent experiments (n =3).

**Table 2.**
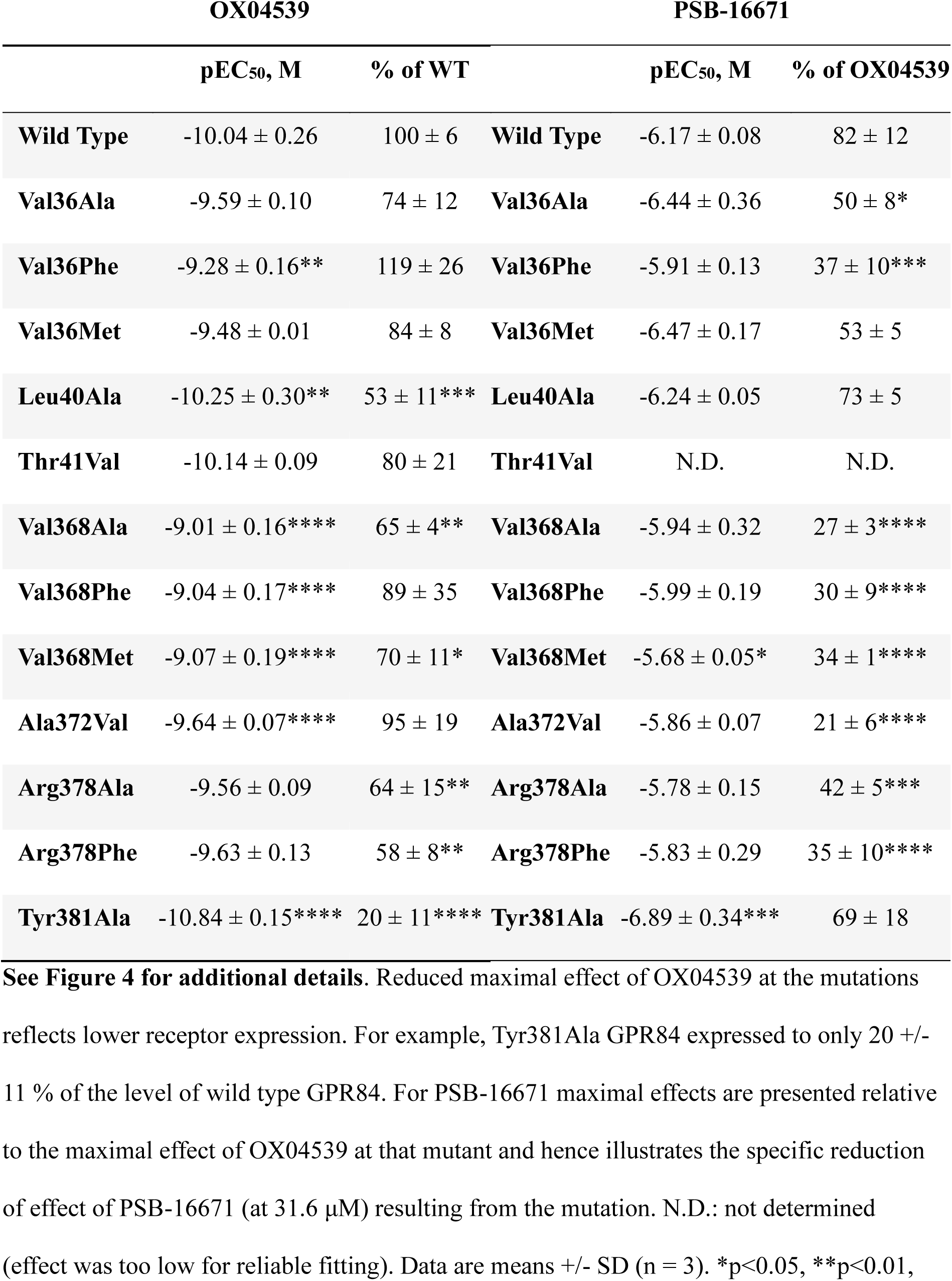
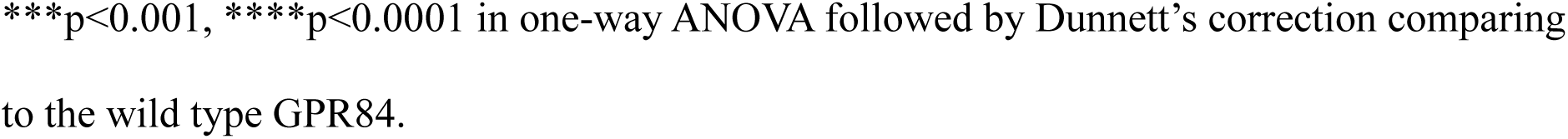
Function of OX04539 and PSB-16671 to directly activate G_i_ at wild type and allosteric site mutations of GPR84.

**Scheme 1.**
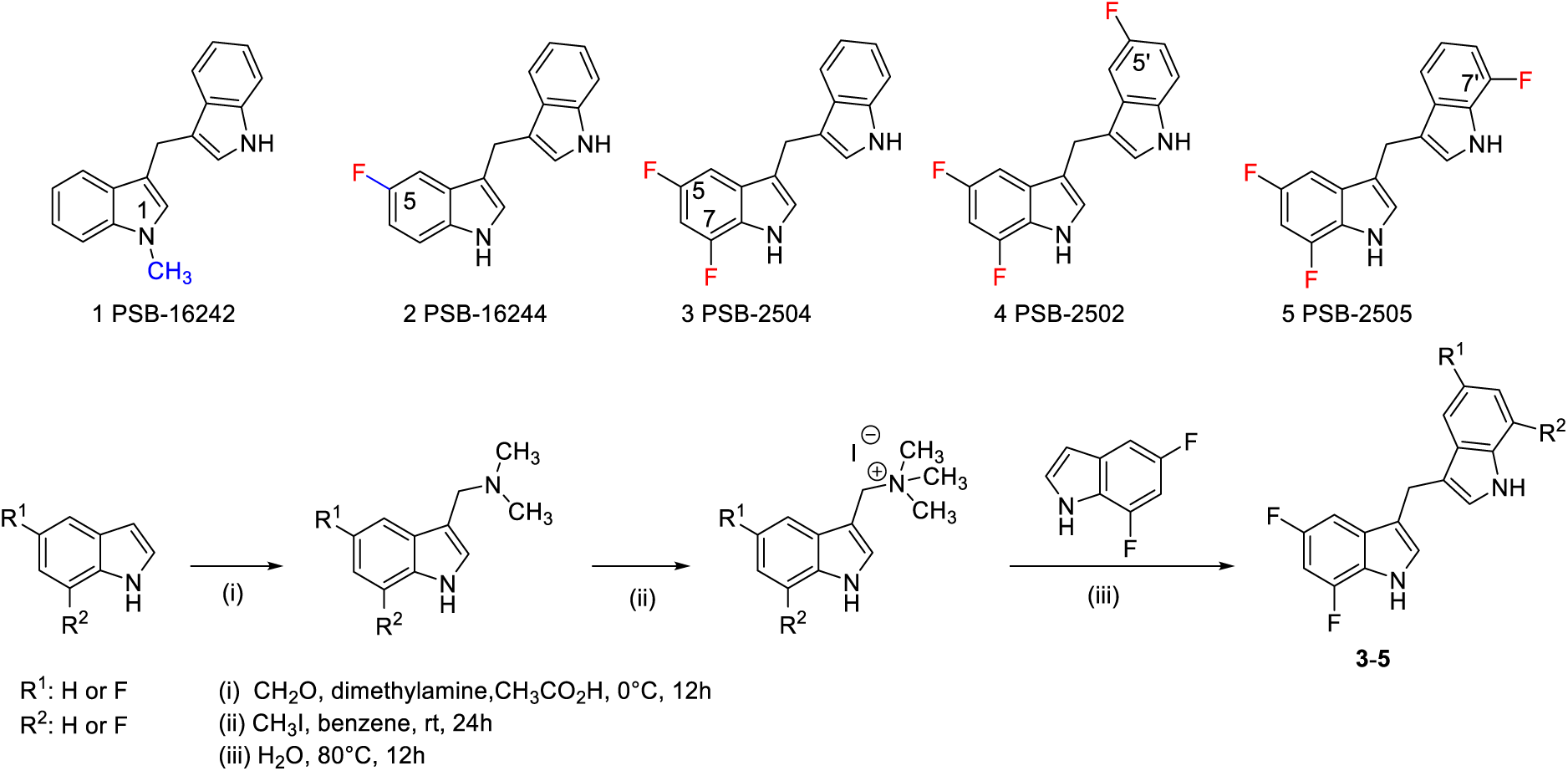
Design and synthesis of additional PSB-16671 derivatives

Additional mutations provided further validation of the binding site. Mutation of Arg378^8.51^ to Ala or to Phe (**Fig. 4g)** reduced response to PSB-16671, consistent with the loss of favorable electrostatic interactions, even when accounting for the observed reduced response to OX04539 that reflects reduced expression of this mutant (**Table 2**). However, although a Tyr381^8.54^Ala alteration appeared to be without major effect on the function of PSB-16671 (**Fig. 4g**), the markedly reduced response observed to OX04539 at this mutant (**Table 2**) indicates a major effect on expression, which made it difficult to explore this alteration in greater detail, whereas mutation of a further potentially important residue, Leu40^1.52^, to Ala was without effect on either expression of this mutant or the function of PSB-16671 (**Fig. 4g**, **Table 2**).

Mutation of the hydrophobic contact residue Val36^1.48^ to Phe, but not to Ala or Met, was detrimental to the PSB-16671 activity (**Fig. 4h**, **Table 2**), while alteration of Val368^7.51^ to Ala, Phe or Met markedly reduced the response to PSB-16671 (**Fig. 4i**, **Table 2**). The poor response of aromatic and Met substitutions at these residues to PSB-16671 indicate they contribute through shape complementarity rather than enhanced interactions with π-electrons of the indole rings.

Notably, for the Ala372^7.55^Val mutation, although G_i_ activation by OX04539 was fully preserved (**Table 2**), the capacity of PSB-16671 to directly activate G_i_ was reduced by ∼83% (mean ± SD, n = 3) (**Fig. 4j**, **Table 2**). Despite this, co-operativity between OX04539 and PSB-16671 still remained at this mutant (**Fig. 4k**). Although there was a trend towards reduced affinity for PSB-16671, this was not statistically significant (p > 0.05) **(Table S1)**. This indicates that while the Ala372^7.55^Val mutation impairs direct G protein activation by PSB-16671, the allosteric communication pathway remains intact.

### OX04539 and PSB-16671 induce distinct active conformations of GPR84

The active conformation of GPR84 in the cryo-EM structures bound to OX04539 alone or to both OX04539 and PSB-16671 closely resembles our previously reported 6-OAU-bound GPR84 structure (**Fig. S4**), which is stabilized by the G_i_ protein. In the orthosteric site, OX04539 is therefore likely to activate the receptor through a mechanism similar to 6-OAU^5^, given their nearly identical binding poses (**Fig. 2b**). This involves the engagement of Tyr332^6.48^ and Phe328^6.44^, which together constitute the ‘transmission switch’ motif critical for GPCR activation^29–32^ (**Fig. S4**).

For PSB-16671, although it sits at the cytoplasmic region of GPR84, it nonetheless induces an active-like conformation of GPR84. To further elucidate the mechanisms by which orthosteric and allosteric ligands regulate GPR84 activation, we performed MD simulations of receptor states with different ligand combinations, in the presence or absence of G_i_ (**Table S3**). Notably, PSB-16671 partially preserves the outward positioning of the cytoplasmic end of TM6, a characteristic feature of GPCR activation, even after the removal of both G_i_ and OX04539 (**Figs. 5a and b**). It also maintains inward conformations of TM7 (**Figs. 5a and c**) and helix 8 (**Fig. S3b**), closely resembling those observed in the ternary complex with G_i_. When combined, OX04539 and PSB-16671 nearly recapitulate the helical structure of G_i_-coupled GPR84, demonstrating their additive effect in stabilizing a G_i_-compatible active conformation of GPR84.

**Fig. 5.**
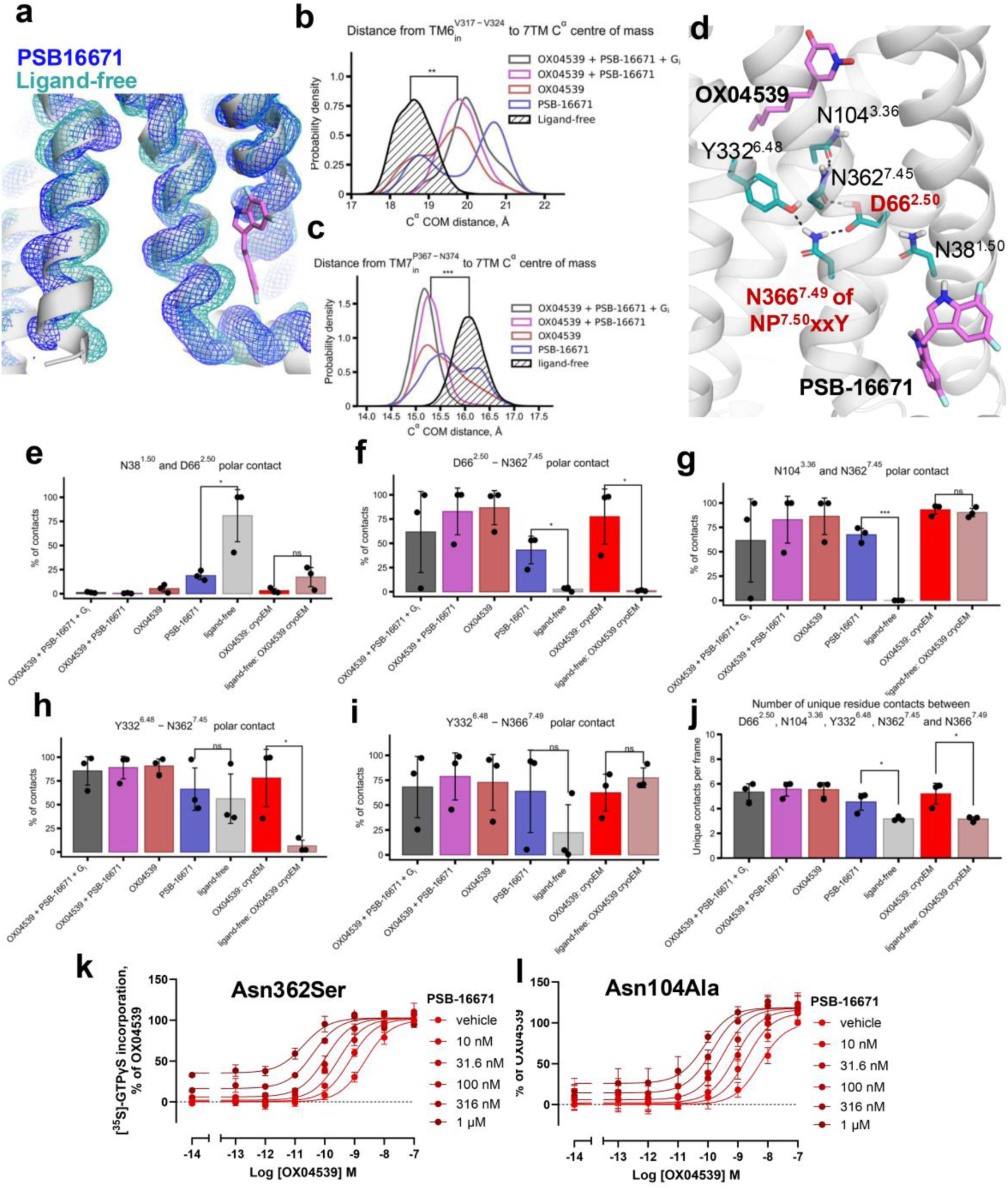
Allosteric biased activation of GPR84 by PSB-16671 via a conserved polar contact network revealed by MD simulations. **a.** Occupancy maps of main chain atoms (N, Cα, C) at 50% iso-value from combined 1 μs simulation replicates (n = 3): PSB-16671-bound (blue) and ligand-free GPR84 (cyan). **b.** Probability density of cytoplasmic TM6 position (Cα centre of mass, Val317^6.33^–Met320^6.36^) measured from 7TM bundle center. Combined frames from three 1 μs replicates per condition. **p < 0.01 (t-test) versus ligand-free. **c.** Probability density of cytoplasmic TM7 position (Cα center of mass, Pro367^7.50^–Asn374^7.57^) measured from 7TM bundle centre. Combined frames from three 1 μs replicates per condition. ***p < 0.001 (t-test) versus ligand-free. **d.** Allosteric communication network in GPR84 showing OX04539 (magenta, orthosteric) and PSB-16671 (magenta, membrane-proximal) with key residues: Asn38^1.50^, Asp66^2.50^ (red), Asn104^3.36^, Tyr332^6.48^, Asn362^7.45^, and Asn366^7.49^ (red, NPxxY motif). Dashed lines indicate hydrogen bonds. **e-i.** Polar contact occupancy (≤4.0 Å) for Asn38^1.50^-Asp66^2.50^, Asp66^2.50^-Asn362^7.45^, Asn104^3.36^-Asn362^7.45^, Tyr332^6.48^-Asn362^7.45^, and Tyr332^6.48^-Asn366^7.49^ sidechains. Combined frames from three 1 μs replicates per condition. *p < 0.05, ***p < 0.001, n.s. = not significant (t-test) versus ligand-free. Error bars: SD between replicates. **j.** Number of polar contacts among Asp66^2.50^, Asn104^3.36^, Tyr332^6.48^, Asn362^7.45^ and Asn366^7.49^. Number of polar contacts is defined as the number of residue-residue pairs forming a polar contact (N/O distance < 4.0 Å) per frame. *p < 0.05 in a simple t-test. Combined frames from three 1 μs replicates per condition. Error bars: SD between replicates. **k-l.** [^35^S]GTPγS incorporation by OX04539 with increasing PSB-16671 concentrations in Asn362^7.45^Ser (**j**) and Asn104^3.36^Ala (**k**) mutants. Data normalized to maximal OX04539 response; mean ± SEM (n=3). Both mutations increase cooperativity despite disrupting polar contacts (**Table 3**).

**Table 3.** Effects on ligand affinity and co-operativity at orthosteric/allosteric site communication residues.

|  | Wild type | Asn104Ala | Asn362Ser |
| --- | --- | --- | --- |
| **logK<sub>(A)</sub>** | -8.83 ± 0.42 | -7.64 ± 0.52* | -7.66 ± 0.17* |
| **logK<sub>(B)</sub>** | -6.16 ± 0.30 | -6.06 ± 0.83 | -5.43 ± 0.14 |
| **logα** | 1.62 ± 0.22 | 1.72 ± 1.14 | 2.75 ± 0.31** |
| **Gain in potency (log)** | 1.26 ± 0.27 | 2.12 ± 0.05* | 2.33 ± 0.24* |

In simulations of the GPR84 Ala372Val mutant, PSB-16671 lacks the ability to stabilize the active-like outward displacement of TM6, whereas OX04539 still induces such feature (**Fig. S3c**), supporting the specific role of Ala372^7.55^ in the ago-allosteric activity of PSB-16671. Consistent with such observation, in our simulations, PSB-16671 forms fewer polar contacts with the Thr41^1.53^ sidechain and the backbone atoms of Thr33^1.45^ and Cys364^7.47^ in GPR84 Ala372Val compared with the wtGPR84, despite remaining within the allosteric pocket (**Fig. S3d**).

Interestingly, PSB-16671 and OX04539 alone induce distinct conformational states of GPR84 in our simulations **(Fig. 5b**). Compared with OX04539, PSB-16671 induces a markedly larger outward displacement of the cytoplasmic end of TM6, even exceeding that observed in the cryo-EM structure stabilized by G_i_ **(Fig. 5b**). Previous structural studies on GPCR-G protein and GPCR-arrestin complexes have revealed that arrestin-coupling is associated with a more modest and smaller outward movement of TM6 relative to G protein coupling^15–17^. The more extensive outward movement of TM6 induced by PSB-16671 is therefore likely incompatible with productive arrestin engagement, providing a structural basis for the G_i_-biased signaling property of PSB-16671.

### Molecular basis for allosteric communication between orthosteric and allosteric sites

In our cryo-EM structure, PSB-16671 is positioned close to Pro367^7.50^ (**Fig. 3b**), a key residue within the conserved NP^7.50^xxY motif in TM7, which undergoes conformational changes during receptor activation^29–32^. We propose that PSB-16671 acts as an ago-PAM by directly modulating the conformation of the NP^7.50^xxY motif, and the allosteric communication between orthosteric and PSB-16671 binding sites operates through a conserved polar contact network involving the NP^7.50^xxY motif and Asp^2.50^ (**Fig. 5d**).

In our MD simulations, both PSB-16671 and OX04539 disrupt the persistent contact between highly conserved residues Asn38^1.50^ and Asp66^2.50^ observed in the ligand-free receptor (**Fig. 5e**), thereby allowing Asp66^2.50^ to engage Asn362^7.45^ (**Fig. 5f**). Asn362^7.45^ subsequently forms hydrogen bonds with the orthosteric site residue Asn104^3.36^ (**Fig. 5g**) and the ’toggle switch’ residue Tyr332^6.48^ (**Figs. 5h and i**). Tyr332^6.48^ in turn interacts with Asn366^7.49^ of the conserved NP^7.50^xxY motif (**Fig. 5i**), establishing a continuous hydrogen-bonding network linking the orthosteric and allosteric sites (**Fig. 5d**). Notably, regardless of the starting cryo-EM structure, the number of contacts among Asp66^2.50^, Asn104^3.36^, Tyr332^6.48^, Asn362^7.45^ and Asn366^7.49^ is higher when either OX04539 or PSB-16671 is present (**Fig. 5j**). This network is partially established by PSB-16671 alone and is fully stabilized when both OX04539 and PSB-16671 are present, demonstrating a cooperative effect between orthosteric and allosteric ligands. This is consistent with our previous work, where we demonstrated that this hydrogen-bonding network plays a critical role in regulating both G protein activation and arrestin recruitment^18^.

Given that residues Asp66^2.50^, Tyr332^6.48^, and Asn366^7.49^ are part of conserved GPCR activation microswitches and their mutations disrupt activation, we generated the GPR84 Asn104Ala and Asn362Ser mutants to explore the functional role of this polar interaction network (**Fig. 5k**). The Asn362Ser mutation significantly reduced both the potency and estimated affinity of OX04539, while only modestly affecting PSB-16671 binding affinity and G_i_ activation (**Fig. 5k, Table S1**). However, surprisingly, this mutation markedly enhanced cooperativity between the two ligands (p < 0.01) (**Table S1)**. Similarly, the Asn104Ala mutation reduced binding affinity of OX04539 without affecting PSB-16671, and once more enhanced their cooperativity (**Fig. 5l, Table S1**).

Simulations of the GPR84 Asn362Ser mutant in the presence of ligands revealed that although a Ser at this position generally retained interactions with Tyr332^6.48^, the frequency of other polar interactions was substantially reduced (**Fig. S3e**). For the Asn104Ala mutation, it disrupts the Asn104^3.36^ and Asn362^7.45^ hydrogen bond interaction, likely increasing the conformational flexibility of Asn362^7.45^ and reducing its interactions with nearby polar residues. Together, these results suggest that disruption of specific polar interactions within this conserved hydrogen-bonding network enhances the cooperativity of orthosteric and allosteric sites by increasing conformational flexibility within the network. This increased flexibility may enable PSB-16671 to more effectively stabilize an active conformation of the orthosteric site that is favourable for agonist binding.

## Discussion

Traditional GPCR drug development has largely focused on orthosteric ligands, but allosteric modulators offer advantages including enhanced subtype selectivity and fine-tuned signaling responses^33–35^. Recent structural studies have revealed diverse allosteric sites across family A GPCRs^36,37^. Here, we identify a helix 8-proximate allosteric site in GPR84 that accommodates the ago-PAM PSB-16671, expanding the structural landscape of GPCR allosteric modulation.

Previously described cytoplasmic PAM sites include the ICL2 region, where PAMs stabilize the E/DRY^3.50^ motif in receptors like D1R^34^, β2AR^35^, and FFA1/2^36,37^, and the G protein-coupling cavity, where PAMs directly engage G proteins in FFA3^38^ and NTSR1^39^ (**Fig. S5**). The helix 8-proximate site in GPR84 represents a third distinct location (**Fig. S5**). While such an allosteric binding pocket is structurally common among GPCRs ^38^, sequence alignments across Class A GPCRs based on the GPCR database (GPCRdb ^39^) show high sequence variations. The overall shape of the pocket is defined by Gly37^1.49^ and to some extent Leu40^1.52^, Val36^1.48^ and Val368^7.51^, which are mostly conserved among class A GPCRs (**Fig. S6**). However, two points of sequence diversity, Thr41^1.49^ and Ala372^7.55^, prevent promiscuous binding of PSB-16671 to the similar allosteric sites in other receptors. Indeed, the prevalent replacement of Thr41^1.49^ with an apolar residue and the even more common replacement of Ala372^7.55^ with a bulky phenylalanine or leucine residues (**Fig. S6**) would disfavor effective binding of PSB-16671 and its analogs to many other Class A GPCRs.

On the other hand, recent preprint findings by Wang et al. indicate that the D1R PAM BMS-A1 ^40^ occupies a similar helix 8-proximate site and forms a critical interaction with Ser36^1.45^, analogous to our Thr41^1.53^-PSB-16671 hydrogen bond. Despite this similar location, the activation mechanisms differ: BMS-A1 disrupts the Thr^1.46^-Ser^7.47^ interaction to maintain D1R’s active state, while PSB-16671 directly modulates GPR84’s conserved NP^7.50^xxY motif in TM7. Our MD simulations and mutagenesis demonstrate that PSB-16671 stabilizes active conformations of TM6, TM7, and helix 8 through an allosteric communication network involving the Asn104^3.36^-Asn362^7.45^-Tyr332^6.48^ hydrogen bond pathway, which can enhance the binding of orthosteric agonists. A similar binding location has also been proposed for a dopamine D2 receptor allosteric modulator based on D2/D3 chimeric studies ^42^. Therefore, while PSB-16771 does not appear to be a broadly promiscuous allosteric modulator of Class A GPCRs, similar helix 8-proximal allosteric sites are likely to exist across this receptor family.

This helix 8-proximate allosteric site in GPR84 also enables biased signaling. PSB-16671 preferentially stimulates G_i_ activation without promoting arrestin recruitment, resulting in sustained phagocytic activity where orthosteric agonists induce desensitization. Such observation highlights the potential of G_i_-biased GPR84 agonism in promoting cancer cell clearance through macrophage-mediated phagocytosis. Mechanistically, our MD simulations suggest that PSB-16671 can induce a receptor conformation characterized by a more pronounced outward displacement of TM6, which is likely incompatible with efficient β-arrestin engagement. Together, our results establish a structural and functional framework for biased agonism at GPR84. The spatial separation of this allosteric site from both the orthosteric ligand-binding pocket and the G protein-coupling interface permits selective modulation of downstream signaling pathways, offering a rational basis for the development of therapeutics with improved signaling profiles.

Our structure validates SAR data showing essential roles for both indole NH groups (hydrogen bonding with Thr41^1.53^ and Thr33^1.45^) and the methylene bridge (occupying a sterically constrained interface)^16,40^. Unsymmetrical analogs with varied fluorination patterns (PSB-2502, PSB-2504, PSB-2505) showed similar potencies, indicating flexibility for future medicinal chemistry optimization. However, selectivity remains a challenge. PSB-16671 retains known off-target activity at CB1, CB2, and P2Y2 receptors^40,41^ and indeed, in functional studies, PSB-16671 has been shown, by an undefined mechanism, to activate G_i_ in mouse bone marrow-derived neutrophils generated from GPR84 knock out mice ^23^. Notably, several critical residues for PSB-16671 binding in GPR84, including Thr41^1.53^, Ala372^7.55^, and F377^8.50^, are not conserved in CB1, CB2, and P2Y2 (**Fig. S6**), arguing against engagement of analogous allosteric sites in these receptors. Consistently, ligand binding experiments using a radioactive orthosteric CB1/CB2 antagonist demonstrated near-complete displacement of the radioligand by DIM ^40^, indicating orthosteric binding of PSB-16671. Nevertheless, the high-resolution structure presented here provides a clear framework for structure-based optimization aimed at improving GPR84 selectivity while preserving beneficial G_i_-biased properties.

## Materials and Methods

### Materials

DIM and 2-HTP (2-(hexylthio)pyrimidine-4,6-diol) was from Merck Life Science UK Ltd. [^35^S]GTPγS, GF/C glassfibre filter-bottom 96-well microplates and Ultima Gold^TM^ XR were from Revvity. [^3^H]140 (3-((5,6-diphenyl-1,2,4-triazin-3-yl)methyl)-1H-indole, 40 Ci/mmol) was produced by Pharmaron (Cardiff, UK) and synthesized as described previously^20^. Compound 837 (3-((5,6-bis(4-methoxyphenyl)-1,2,4-triazin-3-yl)methyl)-1H-indole) was synthesized as described previously^20^. Cell culture reagents, Lipofectamine^TM^ (2000), OptiMem^TM^ and bicinchoninic acid (BCA) assay kit were from Thermo Fisher Scientific. All molecular biology enzymes and reagents were from Promega. Polyethyleneimine (PEI, linear, MW 25,000) was from Polysciences (Warrington, PA). Protease inhibitor mixture was cOmplete^TM^ Mini EDTA-free from Roche Diagnostics.

### Chemical synthesis

The synthesis of several of the investigated DIM derivatives has previously been described ^16,40^. The new unsymmetrical DIM derivatives (PSB-2502, PSB-2504, PSB-2505) were synthesized in analogy to a published procedure^42^ (see Supplementary Data for details).

### Protein expression and purification

The wild-type human GPR84 gene was synthesized and cloned into the pFastBac vector. The construct includes an N-terminal bovine prolactin signal peptide followed by a Flag tag and an 8×His tag. A fragment of β2AR N-terminal tail (BN) was introduced to improve protein expression. To enhance complex stability, the NanoBiT tethering strategy was employed by fusing the LgBiT subunit at the C-terminus of the receptor. The C-terminal residues G388-H396 were truncated, and LgBiT was linked via a 15-amino acid flexible linker (GSSGGGGSGGGGSSG). A dominant negative human Gα_i1_ (DNGα_i1_) containing four mutations (S47N, G203A, E245A, A326S) was cloned into the pFastBac vector. Human Gβ_1_ was fused with an N-terminal His_6_-tag and a C-terminal HiBiT subunit connected with a 15-amino acid linker (GSSGGGGSGGGGSSG), was cloned into pFastBac dual vector together with human Gγ_2_.

ScFv16 was expressed and purified as previously described ^43^. In brief, the protein was expressed in High Five cells and initially purified by nickel affinity chromatography. Further purification was performed using size-exclusion chromatography on a Superdex 200 Increase 10/300 GL column (GE Healthcare). Monomeric peak fractions were pooled, concentrated, and stored at -80 °C before use.

GPR84, DNGα_i1_ and Gβ_1_γ_2_ were co-expressed in Sf9 insect cells using the Bac-to-Bac baculovirus expression system. Cells were co-infected with the three respective baculoviruses at a 1:1:1 ratio. Following incubation at 27 °C for 48 h, cell pellets were harvested and stored at -80 °C until use. Cell pellets were resuspended in lysis buffer containing 20 mM HEPES, pH 7.5, 50 mM NaCl, 10 mM MgCl_2_, 5 mM CaCl_2,_ 2.5 μg/ml leupeptin, 300 μg/ml benzamidine. To facilitate complex formation, 1 μM OX04539, 25 mU/ml Apyrase (NEB), and 100 μM TCEP was added and incubated at room temperature for 2 h. The cell membranes were isolated by centrifugation at 25,000 × g for 30 min and resuspended in solubilization buffer containing 20 mM HEPES, pH7.5, 100 mM NaCl, 0.5% (w/v) lauryl maltose neopentyl glycol (LMNG, Anatrace), 0.1% (w/v) cholesteryl hemisuccinate (CHS, Anatrace), 10% (v/v) glycerol, 10 mM MgCl_2_, 5 mM CaCl_2_, 12.5 mU/ml Apyrase, 1 µM OX04539, 2.5 μg/ml leupeptin, 300 μg/ml benzamidine, 100 µM TECP (tris(2-carboxyethyl)phosphine) for 2h at 4 °C. After centrifugation at 25,000 × g for 45 min at 4 °C, the supernatant was isolated and incubated with Ni resin at 4 °C overnight. The resin was washed with a buffer A containing 20 mM HEPES pH 7.5, 100 mM NaCl, 0.05% (w/v) LMNG, 0.01% (w/v) CHS, 20 mM imidazole, 1 μM OX04539, 2.5 μg/ml leupeptin, 300 μg/ml benzamidine, and 100 μM TCEP. The complex was then eluted using the same buffer supplemented with 400 mM imidazole. The eluate was supplemented with 2 mM CaCl₂ and incubated overnight at 4 °C with anti-Flag M1 antibody resin. The eluate was supplemented with 2mM CaCl_2_ and loaded onto anti-Flag M1 antibody resin. After wash, the complex was eluted in buffer A containing 5 mM EDTA and 200 μg/ml FLAG peptide. Finally, a 1.3 molar excess of purified scFv16 was added to the elution. The sample was then loaded onto a Superdex 200 Increase 10/300 column pre-equilibrated with buffer containing 20 mM HEPES pH 7.5, 100 mM NaCl, 0.00075% (w/v) LMNG, 0.00025% (w/v) GDN, 0.00015% (w/v) CHS, 1 µM OX04539 and 100 µM TECP.

### Cryo-EM sample preparation and data acquisition

The OX04539-GPR84-G_i_ complex was pooled and concentrated to 5 mg/ml for cryo-EM studies. The OX04539-PSB-16671-GPR84-G_i_ complex was prepared by incubating the OX04539-purified complex with 10 µM PSB-16671 for 30 minutes before making the cryo-EM grids. For cryo-EM grids preparation, 3 μl of the purified complexes were applied onto glow-discharged holey carbon grids (Quantifoil, Au300 R1.2/1.3) and plunge-frozen in liquid ethane using a Vitrobot Mark IV (Thermo Fisher Scientific).

For the OX04539-GPR84-G_i_ and OX04539-PSB-16671-GPR84-G_i_ complexes, cryo-EM data were collected on a Titan Krios electron microscope operating at 300 kV, equipped with a Falcon 4i direct electron detector and an energy filter. Micrographs were acquired in super-resolution mode using EPU software at a nominal magnification of 165,000×, yielding a calibrated pixel size of 0.72 Å. The defocus range was set from -1.0 and -2.0 μm. Each image stack was dose-fractionated into 40 frames with a total electron dose of 55 e^-^/Å².

### Data processing, 3D reconstruction and model building

Movies were initially subjected to patch motion correction using cryoSPARC ^44^. Contrast transfer function (CTF) parameters were calculated using the patch CTF estimation tool.

For the OX04539-GPR84-G_i_ complex, a total of 5,605,415 particles were auto-picked and subsequently subjected to 2D classification to eliminate poor-quality particles. After several rounds of 2D classification, the selected particle images were subjected to ab initio reconstruction to generate initial reference maps. Following heterogeneous refinement, 379,621 particles underwent non-uniform refinement and local refinement, resulting in a map with a global resolution of 3.43 Å based on Fourier shell correlation (FSC) at 0.143. Local resolutions of density maps were estimated in cryoSPARC.

For the OX04539-PSB-16671-GPR84-G_i_ complex, a total of 15,430,667 particles were auto-picked and subsequently subjected to 2D classification to eliminate poor-quality particles. After ab initio reconstruction and heterogeneous refinement, 333,622 particles were subjected to non-uniform refinement and local refinement, resulting in a map with a global resolution of 3.22 Å based on Fourier shell correlation (FSC) at 0.143. Local resolutions of density maps were estimated in cryoSPARC.

The model was built based on our previously reported structure of 6-OAU-GPR84-G_i_-scFv16 complex^5^. The structure of GPR84-G_i_-scFv16 was used as initial model for model rebuilding and refinement against the electron microscopy map. The model was docked into the electron microscopy density map using ChimeraX ^45^ followed by iterative manual adjustment and rebuilding in COOT ^46^. Real space refinement was performed using Phenix programs ^47^. The model statistics was validated using programs in Phenix. Structural figures were prepared in ChimeraX and PyMOL ^48^. The final refinement statistics are provided in **Supplementary Table S1-2**.

### Plasmids and mutagenesis

The human GPR84 receptor constructs with an N-terminal FLAG tag and either enhanced yellow fluorescent protein (eYFP), or Gα_i2_-Cys^352^Ile fused to the C-terminus, were cloned into the pcDNA5/FRT/TO expression vector as described previously^19^. Site-directed mutagenesis to generate the point mutations described was performed according to the QuikChange method (Stratagene, Cheshire, UK).

### Cell culture, transfection and generation of cell lines

HEK293T cells were maintained in high-glucose Dulbecco’s modified Eagle’s medium (DMEM) without sodium pyruvate, supplemented with 10% (v/v) fetal bovine serum (FBS), 1% (v/v) penicillin/streptomycin mixture (100 U/ml), and 2 mg/ml normocin (Invivogen), at 37 °C in a 5% CO_2_ humidified atmosphere.

Transient transfection was performed when cells in a 10 cm dish were 60-80% confluent by adding 5 μg of plasmid DNA either in 500 μl of 150 mM NaCl solution with 30 μl PEI or in 500 μl OptiMem^TM^ solution with 15 μl Lipofectamine^TM^.

### [^35^S]GTPγS binding assays

HEK293T cells were transiently transfected with various FLAG-hGPR84-Gα_i2_-C^352^I fusion protein constructs and incubated at 37 °C in a 5% CO_2_ humidified atmosphere. After 24 h, the medium was removed and the cells were suspended in ice-cold phosphate-buffered saline (PBS) and centrifuged at 3000 rpm (1,800 x g) for 5 min at 4 °C with pellets frozen at -80 °C for at least 1 h or longer term. The pellets were resuspended in TE buffer (10 mM Tris, 0.1 mM EDTA, pH 7.4) containing a protease inhibitor mixture and homogenized with a 5 ml hand-held homogenizer, centrifuged at 1200 rpm (450 x g) for 5 min at 4 °C and the supernatant was further centrifuged at 50,000 rpm (90,000 x g, Beckman Ultra Centrifuge) for 45 min at 4 °C. The resulting pellet was resuspended in TE buffer and further homogenized with a 25-gauge needle and protein content was assessed using a BCA protein assay kit.

Prepared membrane protein (5 μg/well, 200 μl reaction volume, 96-well deep-well plates) was incubated in assay buffer (20 mM HEPES, 5 mM MgCl_2_, 160 mM NaCl, 0.05% fatty-acid-free bovine serum albumin, pH 7.4) containing the indicated ligand concentrations. The reaction was initiated by adding [^35^S]GTPγS (100 nCi per reaction) with 1 μM GDP, and the plates were incubated at 30 °C for 60 min. The reaction was terminated by rapid vacuum filtration through GF/C glassfibre filter-bottom 96-well microplates using a UniFilter FilterMate Harvester (PerkinElmer). Unbound radioligand was removed from filters by three washes with ice-cold PBS. Ultima GOLD^TM^ XR was added to dried filters at 50 μl/well, and [^35^S]GTPγS binding was quantified by liquid scintillation spectroscopy.

### Competition binding assays

Competition binding experiments using approximate K_d_ concentrations of [^3^H]140^20^ were performed in binding buffer (phosphate-buffered saline (PBS), 0.5% fatty-acid-free bovine serum albumin; pH 7.4) in a total assay volume of 500 μL in 96 deep-well blocks. Binding was initiated by the addition of 10 μg membrane protein expressing the receptor construct of interest. Nonspecific binding of the radioligand was determined in the presence of 10 μM compound 837. All assays were performed at 25 °C for 60 min before termination by the addition of ice-cold phosphate-buffered saline (PBS) and vacuum filtration through GF/C glassfibre filter-bottom 96-well microplates using a UniFilter FilterMate Harvester (Revvity). Each well was washed three times with ice-cold PBS and filters were allowed to dry for 2–3 h before addition of 50 µl of Ultima Gold^TM^ XR. Radioactivity was quantified by liquid scintillation spectrometry.

### Saturation binding assays

Prepared membrane protein (10 μg/well, 500 μl reaction volume, 96-well deep-well plates) was incubated in phosphate-buffered saline (PBS) with 0.5 % fatty-acid-free bovine serum albumin containing series of [^3^H]140 (40 Ci/mmol) concentrations, with nonspecific binding determined at the same [^3^H]140 concentrations in presence of 10 μM of the GPR84 orthosteric antagonist compound 837^20^. The plates were incubated at 25 °C for 60 min and then harvested by rapid vacuum filtration through GF/C glassfibre filter-bottom 96-well microplates using a UniFilter FilterMate Harvester (PerkinElmer). Unbound radioligand was removed from filters by three washes with ice-cold PBS. Ultima GOLD^TM^ XR was added to dried filters at 50 μl/well, and [^35^S]GTPγS binding was quantified by liquid scintillation spectroscopy. The concentrations of added [^3^H]140 dilutions were quantified by liquid scintillation spectroscopy. To obtain nonspecific binding, from each signal data point the average signal of the respective [^3^H]140 concentration in the presence of 10 μM compound 837 was subtracted.

### Arrestin recruitment Bioluminescence Resonance Energy Transfer (BRET) assays

HEK293T cells were transiently co-transfected with FLAG-hGPR84-eYFP constructs and arrestin-2 (β-arrestin-1) fused to nano-luciferase in a ratio of 100: 1 - 4.95 μg and 0.05 μg, respectively, for each 10 cm dish. As a control, cells were transfected with 100:1 ratio of empty plasmid and nano-luciferase fused arrestin-2. After 24 h, cells were trypsinized and seeded into 96-well white plates coated with poly-D-lysine (40 μl/well, 30 min incubation) at 50,000 cells per well (100 μl/well). The plates were incubated overnight at 37 °C in a 5% CO_2_ humidified atmosphere. After 24 h, the cells were washed once with 100 μl/well Hank’s balanced salt solution (HBSS), pH 7.4, and left in 80 μl/well HBSS for at least 30 min at 37 °C. Then, 10 μl/well of nano-luciferase substrate coelenterazine-h (Nanolight Technologies) was added to a final concentration of 5 μM, and cells were incubated at 37 °C in darkness for 10 minutes. Then, 10 μl/well of orthosteric agonist OX04539 and/or 10 μl/well of allosteric modulator PSB-16671 were added to the plate, and cells were incubated for further 5 min. 15 min after coelenterazine-h addition, reading of emission signals was done on a PHERAstar FS plate reader (BMG Labtech) at 475 nm and 535 nm, which represent nano-luciferase and eYFP emission signals, respectively. The net bioluminescence resonance energy transfer (BRET) ratio (mBRET) was calculated as follows (control is cells without eYFP, see above):

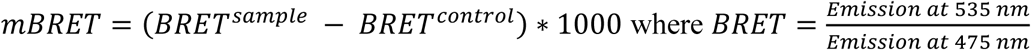

### Experimental data analysis and fitting

Concentration-response curve fitting, one-way ANOVA followed by Dunnett’s multiple comparisons test and simple t-tests were performed using GraphPad Prism version 10.6.1. for Windows, GraphPad Software, Boston, Massachusetts, USA, www.graphpad.com.

Experimental data was normalized with signal at vehicle (no ligands) as 0% and signal at highest OX04539 concentration as 100 %. To compare the OX04539 signal between WT GPR84 and its mutants, the data was normalized to vehicle and OX04539 response of the WT. Unless otherwise stated, experimental graphs represent the Mean ± SEM of pooled average measurements from n = 3 independent biological replicates, and experimental tables represent Mean ± SD of fitted values from n = 3 independent biological replicates.

Concentration-response curves were fitted using three-parameter logistic model:

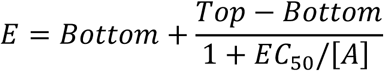

Saturation binding experiments were fitted by one-site specific binding equation to obtain the K_d_ of [^3^H]140:

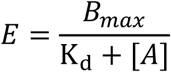

Competition binding experiments were fitted by one-site competition binding equation that employed the measured [^3^H]140 concentration and its K_d_ to obtain the K_i_ of OX04539:

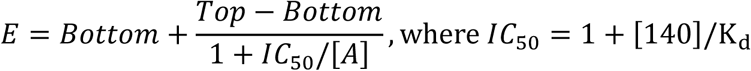

To quantify the extent of allosteric interaction between OX04539 and PSB-16671 in [^35^S]-GTPγS assays, we fitted the signal at varied ligand concentrations to the operational model of allosteric modulation derived^49–52^ and employed^19,53,54^ previously:

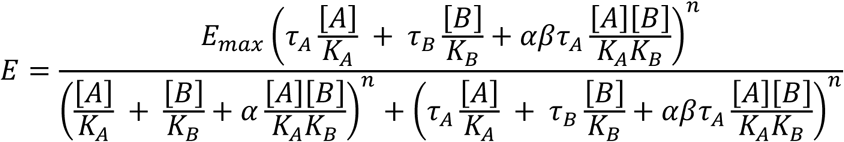

where 𝐸 is the pharmacological effect, [𝐴] and [𝐵] are concentrations of orthosteric ligand (OX04539) and allosteric ligand (PSB-16671), respectively. 𝐾_𝐴_ and 𝐾_𝐵_ are equilibrium dissociation constants of their respective complexes with receptor, 𝛼 is their binding cooperativity factor representing the measure of allosteric effect on orthosteric ligand affinity, 𝛽 is their activation cooperativity factor representing the measure of allosteric effect on orthosteric ligand efficacy, 𝐸 _𝑚𝑎𝑥_ is the maximal possible effect and 𝑛 denotes the slope factor if the transducer function. 𝐸_𝑚𝑎𝑥_ was constrained to the maximum measured response (rounded up) and 𝑛 was constrained to 1, other parameters had no constraints and were estimated by fitting to experimental data. Gain in potency was defined as log 𝐸𝐶_50_^[𝐵]=0^ − log 𝐸𝐶_50_^[𝐵]=1^ ^μM^.

### Cancer cell phagocytosis assay

A luminescence-based long-term phagocytosis assay was performed. Bone marrow–derived macrophages (BMDMs) were generated using mouse bone marrow cells induced by macrophage colony-stimulating factor. BMDMs were pretreated overnight with the GPR84 agonists 6-OAU or PSB16771 (0.0001–10 µM). Following pretreatment, cells were thoroughly washed with PBS to remove residual compounds and subsequently used in the phagocytosis assay. Luciferase-expressing Raji cells were co-cultured with BMDMs for 24 h in the absence or presence of a CD47-blocking antibody (clone B6H12; BioXCell). Luminescence signals were then measured using a Cytation 3 plate reader. Raji cells cultured in the absence of macrophages were used as a normalization control.

### Molecular dynamic simulations

Structures were prepared for simulations and (or) modified with Schrodinger 2025-1 suite ^55–57^. Molecular file format conversions were done with OpenBabel-3.1.0 ^58^, POPC (1-palmitoyl-2-oleoyl-sn-glycero-3-phosphocholine) bilayer systems with 150 mM NaCl were composed with CHARMM-GUI web-server ^59–70^ and simulated with AMBER20 ^71–74^ at 37 °C with OPC water model ^75^ and ff19SB ^76^, lipid21 ^77^ and GAFF2 ^78^ forcefields for protein, lipid and ligand molecules, respectively. Simulations were analyzed via AmberTools24 ^79^ and Python-3.12.11 libraries ^80^, MDAnalysis-2.9.0 ^80^, NumPy-2.3.3 ^81^ and SciPy-1.16.2 ^82^, graphs were built with MatPlotLib-3.10.6 ^83^ in Jupyter Notebooks-4.4.9 ^84^. Occupancy maps were built with Visual Molecular Dynamics (VMD-1.9.4) VolMap tool ^85^, interaction energies were calculated with NAMDEnergy tool employing NAMD2-14 ^86,87^. Structural visualizations were done with opensource PyMOL-3.0.0 ^48^. Detailed methods are given in supplementary information, all simulated systems and their stability parameters are given in Supplementary Table S**3**.

## Acknowledgements

The Pittsburgh Center for CryoEM (RRID:SCR_025216) used for data collection in this project was supported, in part, by the University of Pittsburgh, the School of Medicine, the Department of Structural Biology, and the National Institutes of Health (grants S10-OD-019995 and S10-OD-025009). The content is solely the responsibility of the authors and does not necessarily represent the official views of the National Institutes of Health. This work was supported by the NIH grant R35GM128641 (to C.Z.), the Biotechnology and Biological Sciences Research Council (U.K.) grant BB/T000562/1 (to G.M.), BB/R007101/1 (to I.G.T.), and APP72799 (to AJR, GM and IGT). Support by the Deutsche Forschungsgemeinschaft (DFG) for the Research Training Group GRK 2873 is gratefully acknowledged.

This project made use of computational time on Kelvin-2 supported by Engineering and Physical Sciences Research Council (EPSRC) (grant no. EP/T022175/1 and EP/W03204X/1) and ARCHER2 granted via the UK High-End Computing Consortium for Biomolecular Simulation, HECBioSim (https://www.hecbiosim.ac.uk), supported by EPSRC (grant no. EP/R029407/1 and EP/W03204X/1).

## Author Contributions

CZ, GM, IGT, and AJR designed the study. XZ performed the cryo-EM studies under the supervision of CZ. A-AG, LJ, YL and ZAM performed the computational and pharmacological experiments and analyzed outcomes. JZ performed and analyzed the phagocytosis studies under the supervision of MF. AJR supervised PW who synthesized OX0539. CM supervised FG who synthesized the DIM derivatives and ABM who evaluated their activity at various targets. CZ, GM, IGT, AJR and CM wrote the paper.

## Competing Interests

The authors declare no conflicts of interests.

## Notes

### Competing Interest Statement

The authors have declared no competing interest.

