## Supporting Information for "Mechanism of GPR84 allosteric modulation at a helix 8-proximate site"

### Supplementary Figures

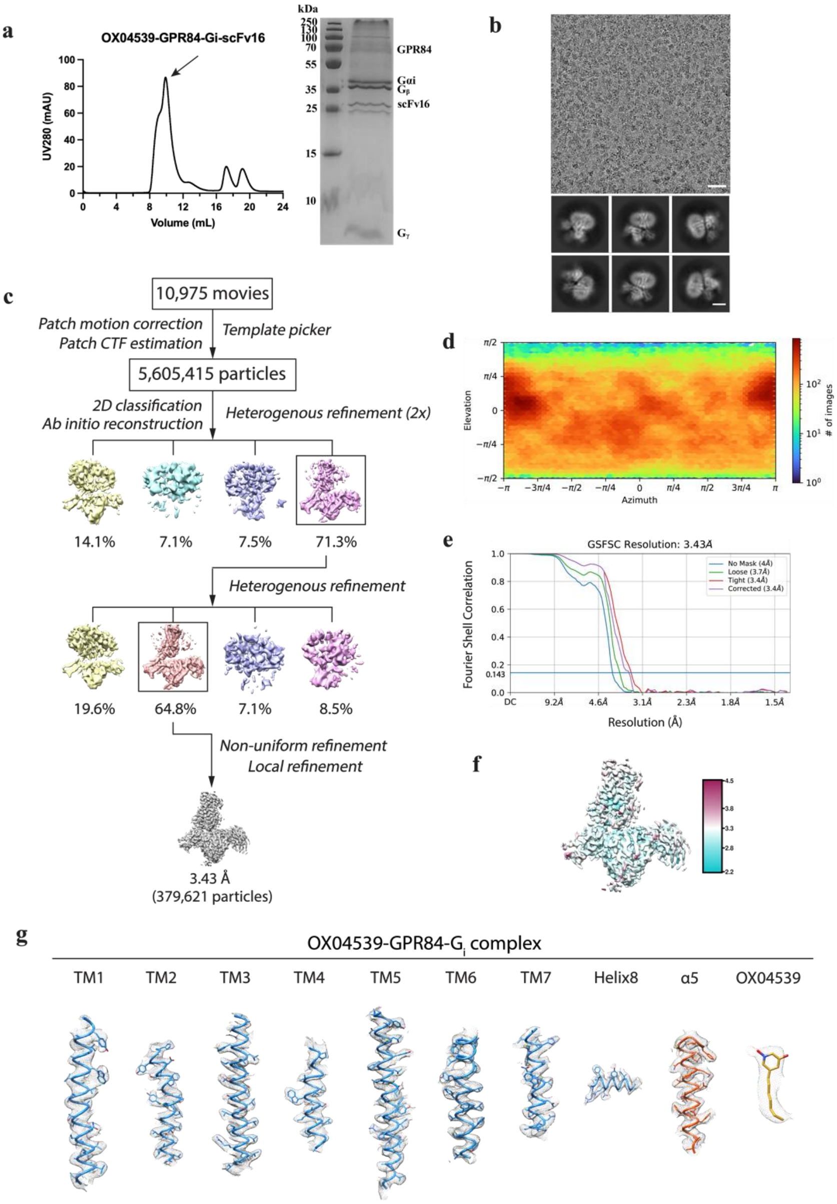


**Supplementary Figure S1. Cryo-EM structure determination of OX04539-GPR84-$G_i$ complex.**

**(a)** Size-exclusion chromatography profile and SDS-PAGE analysis of the purified OX04539-GPR84-$G_i$ complex. **(b)** Representative cryo-EM micrograph (scale bar: 50 nm) and 2D class averages (scale bar: 5 nm). **(c)** Cryo-EM image processing workflow for the OX04539-GPR84-$G_i$ complex. **(d)** Angular distribution of the particles used in the final reconstruction. **(e)** Gold-standard Fourier shell correlation (FSC) curve showing an overall resolution is 3.43 Å at FSC=0.143. **(f)** Density map according to local resolution estimation. **(g)** Cryo-EM density maps and models of the seven transmembrane helices (TM1-7), Helix8, α5 helix of $G_i$ and the ligand of OX04539 bound GPR84-$G_i$ complex are shown.

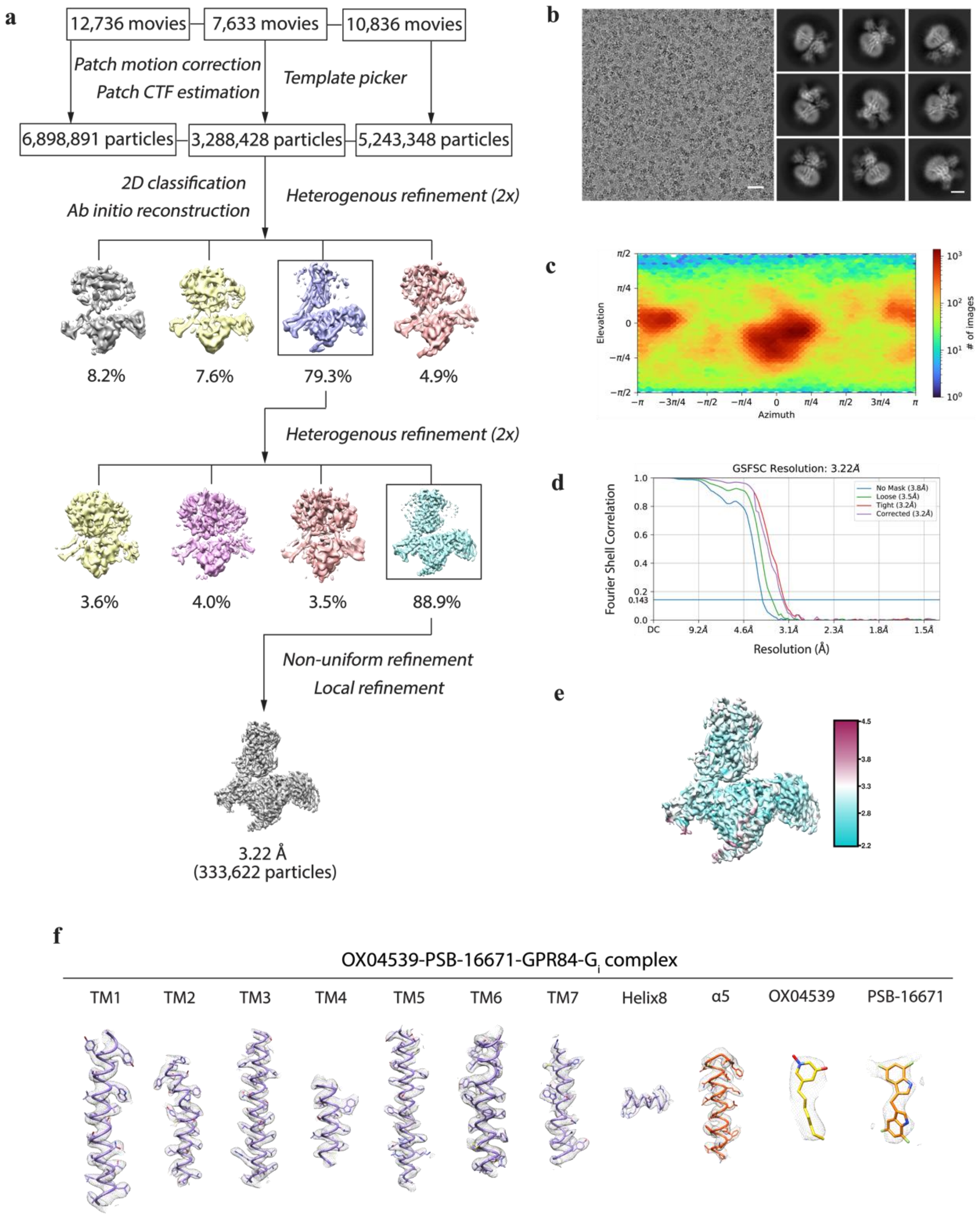


**Supplementary Figure S2. Cryo-EM structure determination of OX04539-GPR84-G$_i$ complex with PSB-16671.**

**(a)** Cryo-EM image processing workflow for the OX04539-PSB-16671-GPR84-G$_i$ complex. **(b)** Representative cryo-EM micrograph (scale bar: 50 nm) and 2D class averages (scale bar: 5 nm). **(c)** Angular distribution of the particles used in the final reconstruction. **(d)** Gold-standard Fourier shell correlation (FSC) curve showing an overall resolution is 3.22 Å at FSC=0.143. **(e)** Density map according to local resolution estimation. **(f)** Cryo-EM density maps and models of the seven transmembrane helices (TM1-7), Helix8, α5 helix of G$_i$ and the ligands of OX04539 and PSB-16671 bound GPR84-G$_i$ complex are shown.

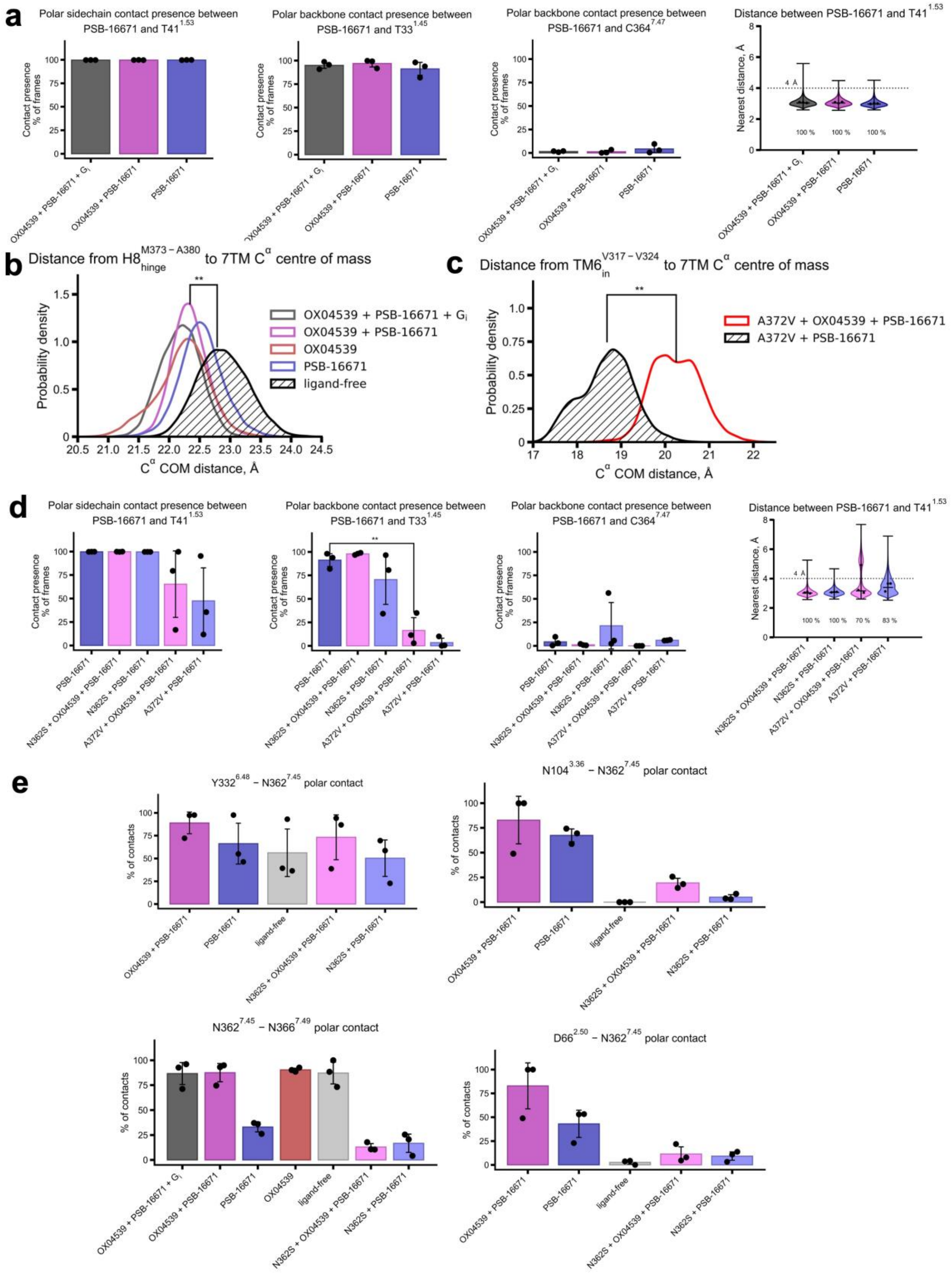


**Supplementary Figure S3. a.** Polar contact occupancy (N/O distance < 4.0 Å) for PSB-16671 and (i) sidechain of Thr41$^{1.53}$, (ii) Thr33$^{1.45}$ or (iii) Cys364$^{7.47}$. Combined frames from three 1 μs replicates per condition. Error bars: SD. (iv) Violin plot showing probability density of PSB-16671 nearest heavy atom distance to Thr41$^{1.53}$ which is located deep within the binding pocket. Combined frames from three 1 μs replicates per condition. Horizontal bars represent minimal, maximal and median distance.

**b.** Probability density of cytoplasmic TM7-H8 region position (Cα center of mass, Met373$^{7.56}$–Ala380$^{8.53}$) measured from 7TM bundle centre. Combined frames from three 1 μs replicas per condition. ** $p<0.01$ (t-test) versus ligand-free. Error bars: SD.

**c.** Probability density of cytoplasmic TM6 position (Cα centre of mass, Val317$^{6.33}$–Val324$^{6.40}$) measured from 7TM bundle center for mutant Ala372Val. Combined frames from three 1 μs replicates per condition. **$p < 0.01$ (t-test) versus OX04539-free.**d.** As for **a.** but for mutants Asn362Ser and Ala372Val in comparison to wild type GPR84.

**e.** Polar contact occupancy (N/O distance < 4.0 Å) for (i) Tyr332$^{6.48}$–Asn362$^{7.45}$, (ii) Asn104$^{3.36}$–Asn362$^{7.45}$, (iii) Asp66$^{2.50}$–Asn362$^{7.45}$ and (iv) Asn362$^{7.45}$–Asn362$^{7.49}$ sidechains. Combined frames from three 1 μs replicas per condition. Error bars: SD.

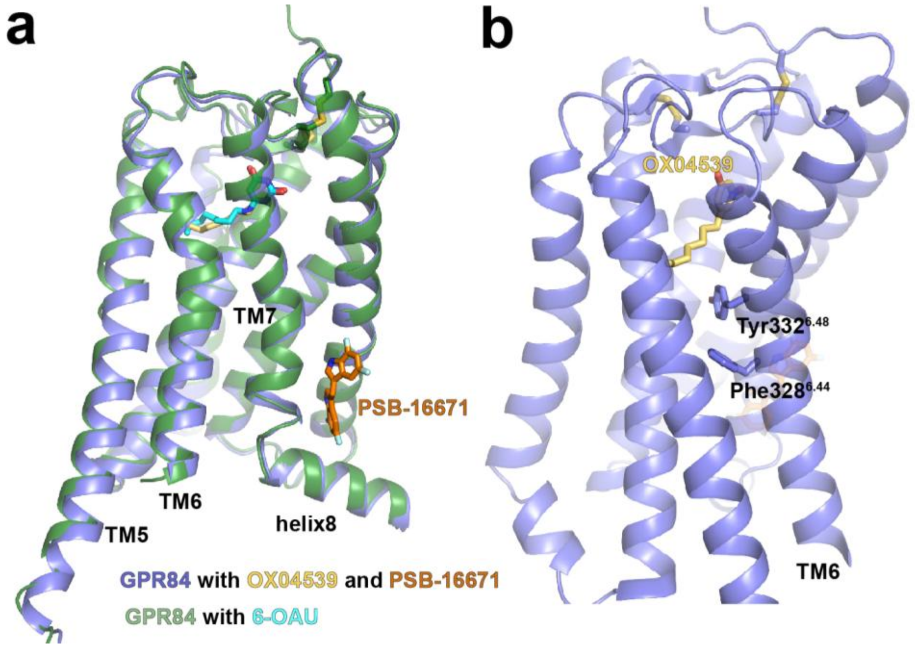


**Supplementary Figure S4. Structure comparison of GPR84 with different ligands. (a)** Superimposition of 6-OAU-bound GPR84 (PDB ID 8G05) and GPR84 bound to OX04539 and PSB-16671. The receptor adopts the same active conformation in both structures stabilized by the Gi protein. **(b)** The orthosteric ligand OX04539 activates the receptor through engagement of the Tyr332$^{6.48}$ and Phe328$^{6.44}$ 'transmission switch' motif in TM6, in a manner similar to 6-OAU.

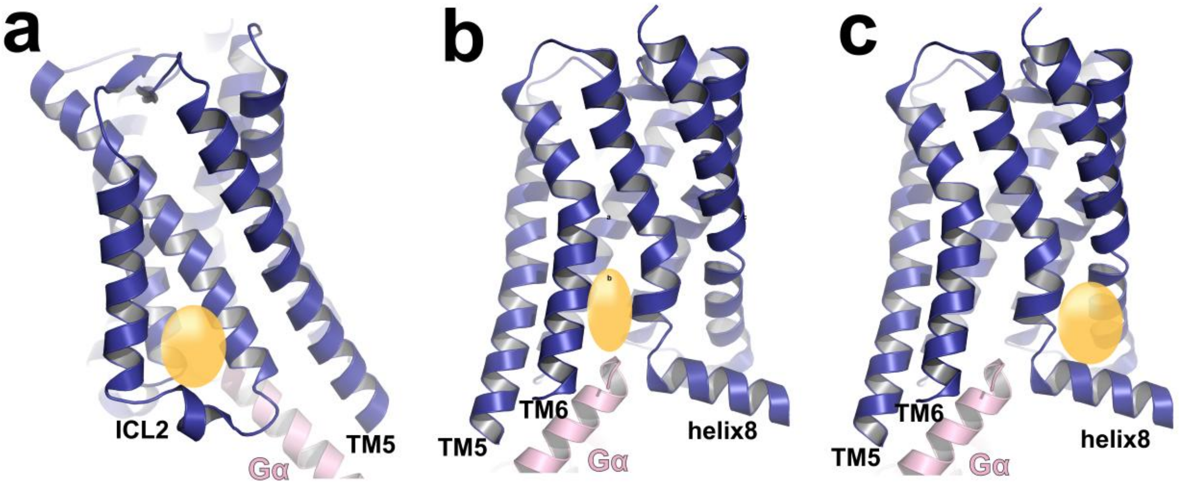


**Supplementary Figure S5. Three representative intracellular allosteric sites in Class A GPCRs for PAMs**. **a.** Site above ICL2. **b.** Site in the G protein-coupled cavity. **c.** Helix 8-proximate site. GPCR and Gα are coloured blue and pink, respectively. Allosteric sites are shown as yellow balls.

**a**

| Residue | Conservation, % | Common residues (conservation in %) |
|---|---|---|
| Thr33 | 4.53 | Val (36 %), Leu (29 %), Ala (9 %) |
| Gly34 | 54.01 | Gly (54 %), Ser (13 %), Ala (11 %) |
| Val36 | 18.82 | Leu (33 %), Val (19 %), Pro (9 %) |
| Gly37 | 67.60 | Gly (68 %), Glu (7 %), Leu (7 %) |
| Leu40 | 56.45 | Leu (56 %), Val (11 %), Ley (6 %) |
| Thr41 | 4.88 | Val (65 %), Ala (14 %), Leu (6%) |
| Cys364 | 39.02 | Cys (39 %), Ala (15 %), Ser (10 %) |
| Ile365 | 15.33 | Leu (34 %), Val (16 %), Ile (15 %) |
| Val368 | 22.30 | Ile (31 %), Val (22 %), Leu (21 %) |
| Ala371 | 25.09 | Ala (25 %), Tyr (10 %), Val (9 %) |
| Ala372 | 2.44 | Phe (43 %), Leu (25 %), Tyr (6 %) |
| Phe377 | 65.51 | Phe (66 %), Leu (7 %), Val (6 %) |
| Arg378 | 54.36 | Arg (54 %), Lys (15 %), Gln (11 %) |
| Tyr381 | 3.83 | Phe (35 %), Leu (30 %), Val (7 %) |

**b**

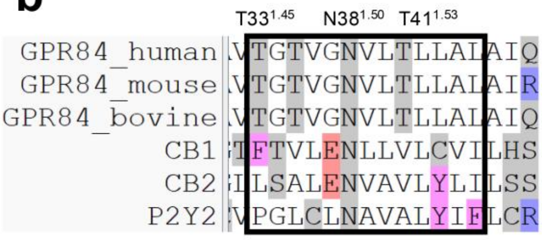


**c**

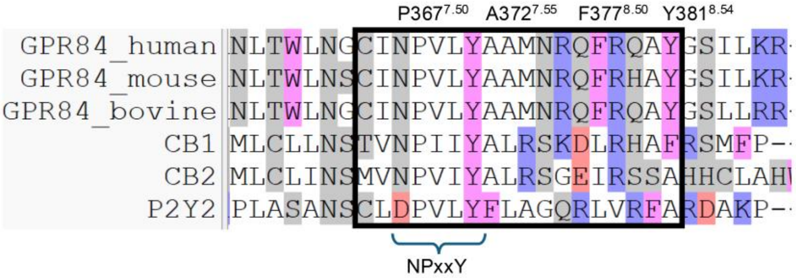


**Supplementary Figure S6. Sequence conservation analysis for residues in PSB-16671 allosteric site across Class A GPCRs**. **a.** Variation of critical residues in the PSB-16671 binding pocket among class A GPCRs based on GPCRdb-based sequence alignments. **b** and **c.** Variation of residues in TM1 (**b**) and TM7-helix 8 (**c**) among human, mouse and bovine GPR84 and human CB1, CB2, and P2Y2 receptors.

### Supplementary Tables

**Table S1. Allosteric co-operativity curve fitting parameters for GPR84 and its mutants**

| | **Wild type** | **Ala372Val** | **Asn104Ala** | **Asn362Ser** |
|---|---|---|---|---|
| **$logK_{(A)}$** | -8.83 ± 0.42 | -8.14 ± 0.41 | -7.64 ± 0.52* | -7.66 ± 0.17* |
| **$logK_{(B)}$** | -6.16 ± 0.30 | -5.80 ± 0.44 | -6.06 ± 0.83 | -5.43 ± 0.14 |
| **logα** | 1.62 ± 0.22 | 2.00 ± 0.78 | 1.72 ± 1.14 | 2.75 ± 0.31** |
| **Gain in potency (log)** | +1.26 ± 0.27 | +1.48 ± 0.45 | 2.12 ± 0.05* | 2.33 ± 0.24* |

Data are extracted from OX04539/PSB-16671 co-operativity studies (see **Fig. 1c** for wild type, **Fig. 4k** for Ala372Val, **Fig. 5k** for Asn104Ala and **Fig. 5l** for Asn362Ser) using the operational model of allosterism. $logK_{(A)}$ is estimated affinity of OX04539, $logK_{(B)}$ is estimated affinity of PSB-16671, α is binding co-operativity factor and gain in potency is the shift in $pEC_{50}$ between the absence and presence of PSB-16671 (1 µM). Data are means +/- SD (n=3). Statistical significance of difference from the wild type was estimated via Welch's t-test.

**Table S2. Cryo-EM data collection and refinement statistics**

| | PW303-GPR84-$G_i$ complex (EMD-75211, PDB ID 10JA) |
|---|---|
| Data collection and processing | |
| Magnification | 165,000 |
| Voltage (kV) | 300 |
| Electron exposure ($e^-/Å^2$) | 50 |
| Defocus range (μm) | -1.0 to -2.0 |
| Pixel size (Å) | 0.72 |
| Symmetry imposed | C1 |
| Initial particle images (no.) | 5,605,415 |
| Final particle images (no.) | 379,621 |
| Map resolution (Å) | 3.43 |
| FSC threshold | 0.143 |
| Map resolution range (Å) | 2.0-4.0 |
| Refinement | |
| Model resolution (Å) | 3.7 |
| FSC threshold | 0.5 |
| Model composition | |
| Non-hydrogen atoms | 7935 |
| Protein residues | 1122 |
| Ligand | 1 |
| R.m.s. deviations | |
| Bond lengths (Å) | 0.003 |
| Bond angles (°) | 0.58 |
| Validation | |
| MolProbity score | 1.78 |
| Clashscore | 9.66 |
| Rotamer outliers (%) | 0.28 |
| Ramachandran plot | |
| Favored (%) | 96.02 |
| Allowed (%) | 3.98 |
| Disallowed (%) | 0 |

| PW303-PSB16671-GPR84-$G_i$ complex (EMD-75212, PDB ID 10JB) | |
|---|---|
| Data collection and processing | |
| Magnification | 165,000 |
| Voltage (kV) | 300 |
| Electron exposure ($e^-/Å^2$) | 50 |
| Defocus range (μm) | -1.0 to -2.0 |
| Pixel size (Å) | 0.72 |
| Symmetry imposed | C1 |
| Initial particle images (no.) | 15,430,667 |
| Final particle images (no.) | 333,622 |
| Map resolution (Å) | 3.22 |
| FSC threshold | 0.143 |
| Map resolution range (Å) | 2.0-4.0 |
| Refinement | |
| Model resolution (Å) | 3.5 |
| FSC threshold | 0.5 |
| Model composition | |
| Non-hydrogen atoms | 8707 |
| Protein residues | 1139 |
| Ligand | 2 |
| Lips:2 clr | |
| R.m.s. deviations | |
| Bond lengths (Å) | 0.004 |
| Bond angles (°) | 0.544 |
| Validation | |
| MolProbity score | 1.66 |
| Clashscore | 9.98 |
| Rotamer outliers (%) | 0.11 |
| Ramachandran plot | |
| Favored (%) | 97.24 |
| Allowed (%) | 2.76 |
| Disallowed (%) | 0 |

**Table S3. RMSDs of simulated structures from the starting structures in MD simulations**

| | RMSD, Å $C^{\alpha}$ | RMSD, Å 7TM $C^{\alpha}$ | RMSD, Å OX04539 | RMSD, Å PSB-16671 | RMSF, Å OX04539 | RMSF, Å PSB-16671 |
|---|---|---|---|---|---|---|
| **OX04539 + PSB-16671 + $G_i$** | 2.15 ± 0.19 | 0.83 ± 0.10 | 0.90 ± 0.04 | 1.78 ± 0.10 | 0.71 ± 0.02 | 0.94 ± 0.01 |
| **OX04539 + PSB-16671** | 2.33 ± 0.23 | 0.84 ± 0.08 | 0.87 ± 0.05 | 1.78 ± 0.03 | 0.71 ± 0.06 | 0.81 ± 0.04 |
| **OX04539** | 2.39 ± 0.35 | 0.83 ± 0.03 | 0.86 ± 0.03 | N.A. | 0.69 ± 0.03 | N.A. |
| **PSB-16671** | 2.53 ± 0.15 | 1.05 ± 0.17 | N.A. | 1.86 ± 0.29 | N.A. | 1.13 ± 0.08 |
| **ligand-free** | 3.15 ± 0.64 | 1.04 ± 0.10 | N.A. | N.A. | N.A. | N.A. |
| **$GPR84^{OX04539}$ + OX04539** | 2.33 ± 0.03 | 0.84 ± 0.11 | 1.04 ± 0.03 | N.A. | 0.70 ± 0.04 | N.A. |
| **$GPR84^{OX04539}$, ligand-free** | 3.37 ± 0.37 | 1.69 ± 0.07 | N.A. | N.A. | N.A. | N.A. |
| **N362S, OX04539 + PSB-16671** | 2.96 ± 0.61 | 0.89 ± 0.02 | 0.93 ± 0.05 | 1.94 ± 0.04 | 0.70 ± 0.05 | 0.83 ± 0.03 |
| **N362S, PSB-16671** | 3.04 ± 0.17 | 1.01 ± 0.08 | N.A. | 1.78 ± 0.22 | N.A. | 1.92 ± 0.11 |
| **A372V, OX04539 + PSB-16671** | 2.73 ± 0.18 | 0.78 ± 0.08 | 0.91 ± 0.07 | 2.75 ± 0.98 | 0.71 ± 0.05 | 2.04 ± 0.68 |
| **A372V, PSB-16671** | 2.38 ± 0.11 | 1.23 ± 0.07 | N.A. | 4.37 ± 1.64 | N.A. | 3.47 ± 0.35 |

Root mean square deviation (RMSD) values are calculated upon alignment by 7TM bundle $C^{\alpha}$ atoms, mean ± SD among averages from 3 replicas of 1 μs simulation. For ligands, only heavy atoms (non-hydrogens) were included in RMSD calculation. N.A.: not applicable (no ligand in simulation). Unless otherwise stated, all simulations were initiated from the GPR84-$G_i$ cryoEM structure in complex with OX04539 and PSB-16671. GPR84$^{OX04539}$ denotes simulations initiated from the GPR84-$G_i$ cryoEM structure in complex with OX04539 alone. 7TM bundle atoms are TM1–TM7 atoms except TM6 below Pro334$^{6.50}$ as described in Supplementary methods.

**Table S4. Calculated residue interaction energies for PSB-16671**

| | Electrostatic energy, kcal.mol$^{-1}$ | Van der Waals energy, kcal.mol$^{-1}$ | Total energy, kcal.mol$^{-1}$ |
|---|---|---|---|
| **T33$^{1.45}$** | -5.29 ± 0.42 | -1.59 ± 0.07 | -6.88 ± 0.07 |
| **L40$^{1.52}$** | 0.27 ± 0.21 | -5.23 ± 0.35 | -4.96 ± 0.35 |
| **T41$^{1.53}$** | -1.88 ± 0.18 | -1.70 ± 0.10 | -3.58 ± 0.10 |
| **G37$^{1.49}$** | -0.19 ± 0.63 | -3.15 ± 0.06 | -3.35 ± 0.06 |
| **C364$^{7.47}$** | -2.18 ± 0.63 | -0.58 ± 0.14 | -2.75 ± 0.14 |
| **V36$^{1.48}$** | -0.01 ± 0.25 | -2.74 ± 0.23 | -2.75 ± 0.23 |
| **F377$^{8.50}$** | -0.98 ± 0.07 | -1.59 ± 0.04 | -2.57 ± 0.04 |
| **R378$^{8.51}$** | -1.04 ± 0.03 | -1.50 ± 0.09 | -2.53 ± 0.09 |
| **Y381$^{8.54}$** | -0.48 ± 0.27 | -1.93 ± 0.58 | -2.41 ± 0.58 |
| **A372$^{7.55}$** | 0.53 ± 0.02 | -2.87 ± 0.10 | -2.33 ± 0.10 |
| **V368$^{7.51}$** | 0.13 ± 0.17 | -1.94 ± 0.08 | -1.81 ± 0.08 |
| **G34$^{1.46}$** | -1.15 ± 0.06 | -0.34 ± 0.01 | -1.49 ± 0.01 |
| **I365$^{1.48}$** | -1.22 ± 0.08 | -0.07 ± 0.02 | -1.28 ± 0.02 |
| **A371$^{7.54}$** | 0.60 ± 0.25 | -1.79 ± 0.45 | -1.19 ± 0.45 |

Molecular mechanics (electrostatic and van der Waals) interaction energies of individual residues with PSB-16671 are calculated for every 10$^{th}$ frame (1 ns interval) of simulation, Mean ± SD among averages from 3 replicas of 1 μs simulation. See **Fig. 3d** for structural visualisation of interaction energies.

**Supplementary Methods**

**Chemical Synthesis**

5-Fluoro-, 7-fluoro- or unsubstituted indole was subjected to a Mannich reaction with formaldehyde and dimethylamine in the presence of acetic acid at 0°C for 12h to yield the corresponding 3-(dimethylaminomethyl)indoles, which were methylated with methyl iodide to the trimethylammonium iodides. Subsequent reaction with 3,7-difluoroindole in water at 80°C provided the desired derivatives in analogy to a published procedure (**Scheme 1**).

Dimethylamine (2M in tetrahydrofuran), formaldehyde (37% w/v in $H_2O$) (1 eq.) and glacial acetic acid (2.3 eq.) were mixed and stirred at 0°C for 10 min. Then, a solution of the appropriate (un)substituted indole derivate (1 eq.) dissolved in glacial acetic acid was dropwise added. The reaction mixture was allowed to warm to room temperature and stirred overnight. After completion of the reaction, the solution was poured into water and brought to an alkaline pH of about 10 using 2N NaOH. The formed precipitate was filtered off and purified by column (DCM/MeOH 9:1) (General Procedure A).

The product was subsequently dissolved in benzene, and methyl iodide (2 eq.) was added. The reaction mixture was then stirred at room temperature overnight. Diethylether was subsequently added, and the precipitate filtered off and washed with diethyl ether. The product was used without further purification and analysis (General Procedure B).

In the final step, the indolylmethylammonium iodide and 5,7-difluoroindole (2 eq.) were suspended in water, and heated in a pressure tube at 80°C overnight upon stirring. After completion of the reaction, the mixture was allowed to cool to room temperature. Then, the water was decanted and the crude solution of the desired product was purified by column chromatography on silica gel with cyclohexane/ethyl acetate (8:2) (General Procedure C).

1-(7-Fluoro-1*H*-indol-3-yl)-*N,N*-dimethylmethanamine was synthesized according to General Procedure A. 7-Fluoro-1*H*-indole (7.45 mmol) was used for the synthesis. A white solid was isolated. Yield 75%.$^{1}$H NMR (500 MHz, DMSO-$d_6$) δ (ppm) 11.4 (s, 1H), 7.4 – 7.4 (m, 1H), 7.3 (s, 1H), 7.0 – 6.8 (m, 2H), 3.5 (s, 2H), 2.1 (s, 6H).$^{13}$C NMR (126 MHz, DMSO-$d_6$) δ (ppm) 149.3 (d, *J* = 242.4 Hz), 131.7 (d, *J* = 5.8 Hz), 125.6, 124.2 (d, *J* = 12.9 Hz), 118.7 (d, *J* = 6.5 Hz), 115.5 (d, *J* = 4.2 Hz), 113.0, 105.8 (d, *J* = 16.1 Hz), 54.4, 45.0.$^{19}$F NMR (471 MHz, DMSO-$d_6$) δ (ppm) -134.8. Purity by HPLC-UV (220-600 nm) ESI-MS: 98.33%, LC-MS calcd. for $C_{11}H_{13}FN_2^-$[M-H]$^-$ 191.11, found 191.10.

5,7-Difluoro-3-((5-fluoro-1*H*-indol-3-yl)methyl)-1*H*-indole was synthesized according to general procedure C. (5-Fluoro-3-indolylmethyl)trimethylammonium iodide (1.62 mmol) and 5,7-difluoro-1*H*-indole (3.27 mmol) were used for the synthesis. A beige-colored resin was isolated. Yield 33%.$^{1}$H NMR (600 MHz, DMSO-$d_6$) δ (ppm) 11.4 – 11.3 (m, 1H, N<u>H</u>), 10.9 – 10.8 (m, 1H, N<u>H</u>), 7.4 (d, *J* = 2.4 Hz, 1H), 7.3 – 7.3 (m, 2H), 7.2 (dd, *J* = 10.1, 2.6 Hz, 1H), 7.1 (dd, *J* = 9.5, 2.2 Hz, 1H), 6.9 – 6.8 (m, 2H), 4.1 (s, 2H, C<u>H$_2$</u>).$^{13}$C NMR (126 MHz, DMSO-$d_6$) δ (ppm) 156.5 (d, *J* = 230.6 Hz), 155.3 (dd, *J* = 233.8 Hz), 148.1 (dd, *J* = 246.1, 14.7 Hz), 133.0, 129.8 (dd, *J* = 11.3, 7.2 Hz), 127.2 (d, *J* = 9.4 Hz), 125.9, 125.0, 120.8 (d, *J* = 12.8 Hz), 115.6 – 115.5 (m), 113.8 (d, *J* = 4.5 Hz), 112.2 (d, *J* = 9.5 Hz), 108.8 (d, *J* = 26.2 Hz), 103.2, 103.1, 99.6 (dd, *J* = 22.9, 3.5 Hz), 96.0 – 95.5 (m), 20.5.$^{19}$F NMR (471 MHz, DMSO-$d_6$) δ (ppm) -124.0 (t, *J* = 9.9 Hz), -126.5 – -126.7 (m), -131.1 (d, *J* = 11.8 Hz). Purity by HPLC-UV (220-600 nm) ESI-MS: 97.86%, LC-MS calcd. for $C_{17}H_{11}F_3N_2^-$[M-H]$^-$ 299.09, found 299.10.

3-((1*H*-Indol-3-yl)methyl)-5,7-difluoro-1*H*-indole was synthesized according to general procedure C. (3-Indolylmethyl)trimethylammonium iodide (1.63 mmol) and 5,7-difluoro-1*H*-indole (3.27 mmol) were used for the synthesis. A beige-colored resin was isolated. Yield 40%.$^{1}$H NMR (500 MHz, DMSO-$d_6$) δ (ppm) 11.4 – 11.3 (m, 1H, N<u>H</u>), 10.7 (s, 1H, N<u>H</u>), 7.5 (dd, *J* = 7.8, 1.1 Hz, 1H), 7.3 – 7.3 (m, 2H), 7.2 (d, *J* = 2.3 Hz, 1H), 7.1 (dd, *J* = 9.6, 2.3 Hz,

1H), 7.0 (m, $J$ = 8.2, 7.0, 1.2 Hz, 1H), 6.9 – 6.8 (m, 2H), 4.1 (s, 2H, C<u>H</u>$_2$).$^{13}$C NMR (151 MHz, DMSO-d$_6$) δ (ppm) 155.1 (dd), 149.6 – 146.6 (m), 136.5, 130.1 – 129.7 (m), 127.1, 125.9, 123.0, 120.9, 118.7, 118.2, 116.3 – 115.7 (m), 113.6, 111.4, 100.1 – 99.3 (m), 95.9 (dd, $J$ = 30.5, 21.0 Hz), 59.9, 20.8.$^{19}$F NMR (471 MHz, DMSO-d$_6$) δ (ppm) -124.1 (t, $J$ = 10.2 Hz), -131.2 (d, $J$ = 12.1 Hz). Purity by HPLC-UV (220-600 nm) ESI-MS: 95.56%, LC-MS calcd. for $C_{17}H_{12}F_2N_2^-$[M-H]$^-$ 281.10, found 281.10.

5,7-Difluoro-3-((7-fluoro-1*H*-indol-3-yl)methyl)-1*H*-indole was synthesized according to general procedure C. (7-Fluoro-3-indolylmethyl)trimethylammonium iodide (1.63 mmol) and 5,7-difluoro-1*H*-indole (3.27 mmol) were used for the synthesis. A beige-colored resin was isolated. Yield 41% .$^1$H NMR (500 MHz, DMSO-d$_6$) δ (ppm) 11.4 – 11.3 (m, 1H, N<u>H</u>), 11.3 – 11.2 (m, 1H, N<u>H</u>), 7.4 – 7.3 (m, 1H), 7.3 (d, J = 2.3 Hz, 1H), 7.3 (d, J = 2.3 Hz, 1H), 7.1 (dd, J = 9.6, 2.2 Hz, 1H), 6.9 – 6.8 (m, 3H), 4.1 (s, 2H, C<u>H</u>$_2$).$^{13}$C NMR (126 MHz, DMSO-d$_6$) δ (ppm) 155.5 (dd, $J$ = 233.5, 9.8 Hz), 149.4 (d, $J$ = 242.4 Hz), 149.3 (d, $J$ = 14.5 Hz), 147.3 (d, $J$ = 14.7 Hz), 131.3 (d, $J$ = 5.9 Hz), 129.9 (dd, $J$ = 11.0, 6.9 Hz), 126.0, 124.3, 124.2 (d), 121.0 (d, $J$ = 12.7 Hz), 118.6 (d, $J$ = 6.1 Hz), 115.7 (dd, $J$ = 5.0 Hz), 115.0 (dd), 105.9, 105.7, 99.8 (dd, $J$ = 22.8, 3.7 Hz), 96.0 (dd, $J$ = 30.5, 21.0 Hz), 20.8.$^{19}$F NMR (471 MHz, DMSO-d$_6$) δ (ppm) -124.0 (t, $J$ = 9.5 Hz), -131.1 (d, $J$ = 10.5 Hz), -134.8 (dd, $J$ = 12.0, 5.1 Hz). Purity by HPLC-UV (220-600 nm) ESI-MS: 98.47%, LC-MS calcd. for $C_{17}H_{11}F_3N_2^-$[M-H]$^-$ 299.09, found 299.30.

***Classical MD simulations***

MD simulations were conducted on GPR84 systems derived from the two cryo-EM complexes obtained in this work by removing individual components and (or) introducing mutations. The structures were prepared for simulations using Schrodinger Maestro 2025-1[2], missing sidechains and loops were filled up to 10 residues (up to 5 from each end of the sequence) as per the UniProt[40] canonical sequence via Schrodinger Prime[1,3,4] (Homology Model Building protocol), clashes between parts of the system were relaxed by local energy minimization in 3D Builder utility in OPLS4 forcefield[41,42], mutations were also introduced via 3D Builder. All histidine residues were taken as δ-tautomers, Asp66[2.50] was protonated as per PROPKA3 prediction[43,44], other titratable residues were taken at their protonation state under pH 7 (positive lysine and arginine residues, negative aspartate and glutamate residues).

Membrane bilayer systems for simulations were composed with CHARMM-GUI server[6–17], with protein oriented via their OPM server [45,46] communication tool and placed in system with a 100 Å x 100 Å membrane (when G protein is bound, 130 Å x 130 Å) containing the automatically estimated number of 1-palmitoyl-2-oleoyl-sn-glycero-3-phosphocholine (POPC) and with a 22.5 Å layer of water on each side, water contained 150 mM NaCl on top of charge-balancing ions, making a total of 100,000 – 130,000 atoms (200,000 – 250,000 for G-protein bound simulation).

Simulations of Langevin dynamics at the pressure of 1 bar and the temperature of 310 K (37 °C) were conducted in AMBER20[21] with GPU implementation[18–20]. We used OPC water model[22] and ff19SB[23], lipid21[24] and GAFF2[25] forcefields for protein, lipids and ligands, respectively, long range interaction cutoff was 9 Å, holonomic constraints were SHAKE[47]. The Langevin thermostat[48,49] had a friction coefficient of 1 $ps^{-1}$, and the pressure control where needed was done by Berendsen barostat[50] with $\tau_{couple}$ of 1 $ps^{-1}$.

Minimization and equilibration followed a modification of default CHARMM-GUI protocol. Minimization was done with 15,000 steepest descend steps and 35,000 conjugated gradient steps, positional restraints were applied to protein and ligand atoms (10.0 kcal/mol/$Å^2$) and lipid P-atoms (2.5 kcal/mol/$Å^2$). Velocities were randomly generated at target temperature with SEED taken from machine time in microseconds. Equilibration was done in 7 steps with gradually diminishing restraints: 2 steps of 2.5 ns NVT simulation with 1 fs timestep, 1 step of 2.5 ns NPT simulation with 1 fs timestep, 2 steps of 10 ns with 2 fs timestep and 2 steps of 20 ns NPT simulation. The positional restraints on protein and ligand atoms at steps 1 – 7 relaxed as 10.0, 5.0, 2.5, 1.0, 0.5, 0.1 and 0.0 kcal/mol/Å$^2$, with an additional constraint on distance between PSB-16671 polar hydrogen and oxygen of Cys364$^{7.47}$ that appeared when their distance exceeded 4 Å and relaxed as 10.0, 10.0, 5.0, 2.5, 1.0, 0.5 and 0.5 kcal/mol/Å$^2$. The positional restraints on lipid P-atoms relaxed as 2.5, 2.5, 1.0, 0.5, 0.1, 0.0 and 0.0 kcal/mol/Å$^2$, and dihedral restraints relaxed as 250, 100, 50, 50, 25, 0, 0 kcal/mol/°$^2$. Production runs in 3 replicates of 1,000 ns (1 µs) simulation were initiated from the same equilibrated structure, snapshots were taken every 100 ps (0.1 ns).

For analysis, initial structures were aligned to the PPM3-oriented GPR84/6-OAU structure (PDB ID 8G05) by C$^\alpha$ atoms of 7TM bundle, and simulation frames were aligned to the respective initial structure in the same way. Hereafter, 7TM bundle is defined as residues Tyr21-Leu45, Leu58-Leu82, Val95-Tyr119, Ile137-Ser151, Leu182-Tyr198, Pro334-Leu341, Val351-Asn366, thus excluding the flexible TM6 cytosolic portion. Alignment and trajectory-topology processing involved Ambertools24[26,27] CPPTRAJ-6.24[51] utility and MDAnalysis-2.9.0[28,29]. Root Mean Square Deviations (RMSD), distances and contact counts were calculated with python scripts employing MDAnalysis-2.9.0[28,29], NumPy-2.3.3[30] and SciPy-1.16.2[31], graphs were plotted with MatPlotLib-3.10.6[32] in Jupyter Notebooks-4.4.9[33]. Polar contacts were defined as any two polar heavy atoms (N, O, S) being within 4 Å of each other in the given simulation frame. Number of unique polar residue contacts was defined as the number of residue pairs forming at least one polar contact in the given frame. Visual analysis of trajectories involved Visual Molecular Dynamics (VMD-1.9.4)[34]. Occupancy maps were built with VMD-1.9.4[34] VolMap tool on a 0.5 Å grid. Forcefield-level interaction energies (point-charge electrostatic and smoothed 6-12 Lennard-Jones) were calculated with VMD-1.9.4. NAMDEnergy utility employing NAMD-2.14[35,36] with the same forcefield as used for simulation, with the same 9 Å interaction cutoff and smoothing switch distance of 7.5 Å, calculation was done for every $10^{th}$ frame of simulation, i.e. with 1 ns timestep. Structural visualizations were done with opensource PyMOL-3.0.0[37].

26. D.A. Case, H.M. Aktulga, K. Belfon, I.Y. Ben-Shalom, J.T. Berryman, S.R. Brozell, D.S. Cerutti, T.E. Cheatham, III, G.A. Cisneros, V.W.D. Cruzeiro, T.A. Darden, N. Forouzesh, M. Ghazimirsaeed, G. Giambaşu, T. Giese, M.K. Gilson, H. Gohlke, A.W. Goetz, J. Harris, Z. Huang, S. Izadi, S.A. Izmailov, K. Kasavajhala, M.C. Kaymak, A. Kovalenko, T. Kurtzman, T.S. Lee, P. Li, Z. Li, C. Lin, J. Liu, T. Luchko, R. Luo, M. Machado, M. Manathunga, K.M. Merz, Y. Miao, O. Mikhailovskii, G. Monard, H. Nguyen, K.A. O'Hearn,

A. Onufriev, F. Pan, S. Pantano, A. Rahnamoun, D.R. Roe, A. Roitberg, C. Sagui, S. Schott-Verdugo, A. Shajan, J. Shen, C.L. Simmerling, N.R. Skrynnikov, J. Smith, J. Swails, R.C. Walker, J. Wang, J. Wang, X. Wu, Y. Wu, Y. Xiong, Y. Xue, D.M. York, C. Zhao, Q. Zhu, and P.A. Kollman (2024), Amber 2024, University of California, San Francisco. (2024).

37. Schrodinger, LLC. 2010. The PyMOL Molecular Graphics System, Version 3.0.
